# Transcriptomic analysis reveals a role for the nervous system in regulating growth and development of *Fasciola hepatica* juveniles

**DOI:** 10.1101/2022.01.13.476286

**Authors:** Emily Robb, Erin McCammick, Duncan Wells, Paul McVeigh, Erica Gardiner, Rebecca Armstrong, Paul McCusker, Angela Mousley, Nathan Clarke, Nikki Marks, Aaron G. Maule

## Abstract

*Fasciola* spp. liver fluke have significant impacts in veterinary and human medicine. The absence of a vaccine and increasing anthelmintic resistance threaten sustainable control and underscore the need for novel flukicides. Functional genomic approaches underpinned by *in vitro* culture of juvenile *Fasciola hepatica* facilitate control target validation in the most pathogenic life stage. Comparative transcriptomics of *in vitro* and *in vivo* maintained 21 day old *F. hepatica* finds that 86% of genes are expressed at similar levels across maintenance treatments suggesting commonality in core biological functioning within these juveniles. Phenotypic comparisons revealed higher cell proliferation and growth rates in the *in vivo* juveniles compared to their *in vitro* counterparts. These phenotypic differences were consistent with the upregulation of neoblast-like stem cell and cell-cycle associated genes in *in vivo* maintained worms. The more rapid growth/development of *in vivo* juveniles was further evidenced by a switch in cathepsin protease expression profiles, dominated by cathepsin B in *in vitro* juveniles and then by cathepsin L in *in vivo* juveniles. Coincident with more rapid growth/development was the marked downregulation of both classical and peptidergic neuronal signalling components in *in vivo* maintained juveniles, supporting a role for the nervous system in regulating liver fluke growth and development. Differences in the miRNA complements of *in vivo* and *in vitro* juveniles identified 31 differentially expressed miRNAs, notably *fhe-let-7a-5p*, *fhe-mir-124-3p* and, miRNAs predicted to target Wnt-signalling, supporting a key role for miRNAs in driving the growth/developmental differences in the *in vitro* and *in vivo* maintained juvenile liver fluke. Widespread differences in the expression of neuronal genes in juvenile fluke grown *in vitro* and *in vivo* expose significant interplay between neuronal signalling and the rate of growth/development, encouraging consideration of neuronal targets in efforts to dysregulate growth/development for parasite control.

**Author Summary:** Parasitic worms are notoriously difficult to study outside of a host organism. However, recent developments in culture methods for *Fasciola hepatica* liver fluke juveniles support growth and development of these parasites in the laboratory (*in vitro*) towards adult parasites. Having the ability to grow pathogenic juvenile stages *in vitro* enables functional studies to validate potential drug and vaccine targets. However, comparison of *in vitro* grown juveniles to juveniles retrieved from infected hosts (*in vivo*) shows considerable size differences suggesting at least some differences in biology that could undermine the relevance of data generated from *in vitro* maintained parasites. This study examines gene expression differences between *in vitro* and *in vivo* maintained *F. hepatica* juveniles via transcriptomic analysis to identify similarities and differences in their biology which may help explain differences in the rate of growth and development. 86% of genes were shown to be expressed at similar levels across treatment groups suggesting a high level of biological similarity between *in vitro and in vivo* juveniles. However, the genes that are expressed differently between these juveniles will help improve current culture methods and provide a new group of potential drug targets that impact on juvenile growth and development.

## Introduction

*Fasciola spp*. liver fluke are important helminth pathogens with far reaching global impacts on veterinary and human medicine causing a disease known as fasciolosis [1]. Global agricultural losses associated with *Fasciola* infection are estimated at around US$3.2 billion annually [2], although this is thought to be a considerable underestimation as *F. hepatica* parasites in particular are known to infect a wide range of mammalian hosts across broad geographical ranges [3, 4]. Designated a neglected tropical disease by the World Health Organisation (WHO) [1] due to its impact on human populations, *F. hepatica* is thought to infect up to 17 million people, with a further 91 million people at risk of infection worldwide [5].

Treatment of the early pathogenic stages of fasciolosis relies on the drug Triclabendazole, since other flukicides cannot kill young juveniles [4, 6]. Definitive mammalian hosts are infected through the ingestion of infectious metacercariae encysted on vegetation [7, 8]. Newly excysted juveniles (NEJs) emerge in the duodenum and migrate through the intestinal wall to the liver parenchyma, causing acute stages of disease associated with host blood loss and even death with high burden infections [7–9]. Although adults residing in the bile duct are reproductively active and associated with chronic infection, the major pathology associated with fasciolosis is caused by juvenile migration within the first few weeks of infection [10, 11]. The absence of a liver fluke vaccine and increasing anthelmintic resistance threaten the sustainability of liver fluke control. Triclabendazole treatment failure is widely reported for livestock and more recent cases of emerging resistance in human populations is deeply concerning and highlights the pressing need for new flukicides, particularly targeting the highly pathogenic early-stage juveniles [4, 12–14].

Despite the pathology associated with early-stage infections, there is a dearth of information on the biology of migrating juveniles due to their small size, liver parenchyma location and, historically, the absence of readily amenable *in vitro* culture methods [15]. At this stage of development, juveniles are growing and moving rapidly through host tissues, encountering different microenvironments and host responses [11]. Understanding the biology of these behaviours and host-parasite interactions will support new target discovery and control option developments. Research into helminth infections often relies on the use of animal models to inform host-parasite relationships. However, such studies are often challenging, of variable relevance to livestock/human hosts and provide limited opportunity to interrogate parasite biology [16]. *In vitro* culture, where possible, supports curiosity driven research and provides many advantages to early-stage studies of parasite biology. However, despite recent advances in propagation of the full life cycle of *Schistosoma mansoni* [17], *in vitro* parasite culture is notoriously difficult and, where it is possible, the approaches need to reasonably replicate *in vivo* parasite biology such that readouts have relevance to control.

McCusker *et al.* [15] developed a chicken serum-based culture platform that promotes sustained *F. hepatica* juvenile survival, growth and development *in vitro*. This has facilitated the development of a robust functional genomics platform that promotes discovery-based studies that seed drug target identification and validation in the most damaging life stage [18]. Although extremely robust, the current culture platform is believed to support a slower rate of growth and development of *F. hepatica* juveniles *in vitro* when compared to *in vivo* counterparts and, whilst *in vitro* juveniles start to develop adult like features, they do not progress to egg laying adults [15]. In the absence of culture methods that allow full life cycle propagation, it is important to ascertain if the slower growing *in vitro* worms resemble their *in vivo* counterparts to ensure functional genomics studies on *in vitro* cultured juveniles have relevance to control.

Taking advantage of recent developments in genomic and transcriptomic resources for *F. hepatica*, we present in depth comparative transcriptomic analysis of 21 day *in vitro* maintained *F. hepatica* juveniles and stage matched 21 day *in vivo F. hepatica* juveniles, describing key differences in gene expression in the most pathogenic life stage under differing growth conditions. Although the expression of core biological functioning components is consistent in comparisons between the two growth conditions, the differences that are evident can mostly be explained by the disparate rates of growth and cell proliferation observed in the two groups. Notably, protease expression profiles differ and corroborate the hypothesis that time-matched *in vitro* maintained juveniles develop more slowly than *in vivo* juveniles, such that when time-matched, the *in vitro* juveniles are at an earlier stage of development. The data indicate a key role for miRNAs in driving the developmental differences between *in vitro* and *in vivo* maintained juveniles with over one third of the identified miRNAs being differentially expressed. Further, profound differences in the expression of genes associated with neoblast proliferation and development are consistent with the observed higher rate of neoblast-like stem cell proliferation *in vivo*. The unexpected down-regulation of multiple nervous system associated signalling pathways exposes an important role for the nervous system of *F. hepatica* in the modulation of juvenile growth and development, encouraging efforts to identify neuronal signalling pathways that dampen cell proliferation and/or growth dynamics.

## Results

A total of 30,892 transcripts were identified from this analysis (S1 Text) which were further refined to define a dataset of 19,343 unique genes for 21 day old juvenile *F. hepatica* (removal of transcript isoforms and non-zero read counts from *in vitro* and *in vivo* assembled transcriptomes) reflective of the greatest depth of sequencing achieved to date for *F. hepatica* (average 252 million mapped reads per replicate transcriptome, S2 Table) [19–23]. All genes were annotated using a core annotation pipeline described in Fig 1. Annotations were assigned to 64% (12,300) of genes based on matches from at least one database interrogated (S3 Table).

**Fig 1:**
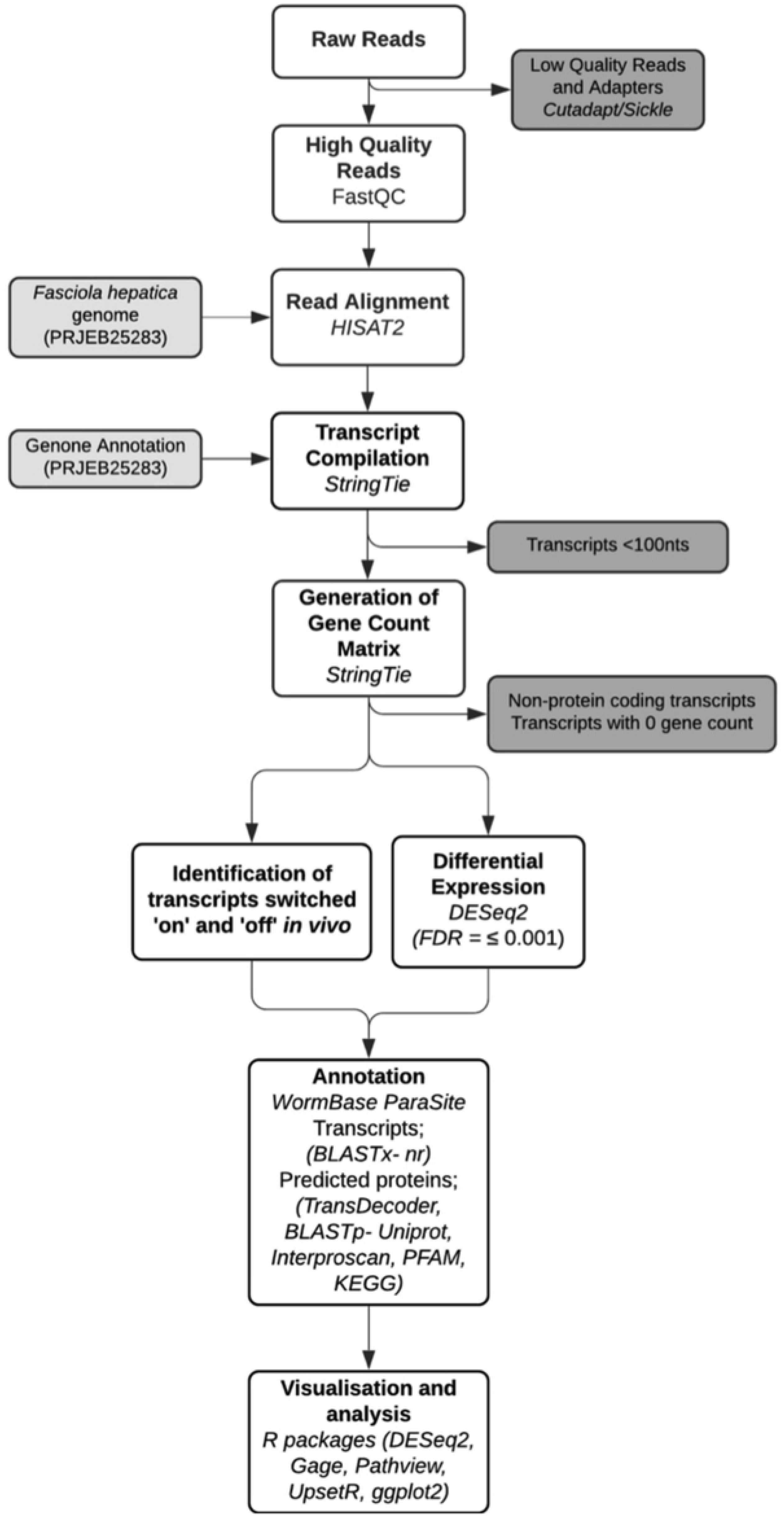
Flow chart of bioinformatics pipeline. Outline of methods used to assemble and analyse transcriptomes. Low quality reads and adapters were removed using Cutadapt (v.1.2.1) and Sickle (v.1.200) and remaining reads quality assessed using FastQC (v. 0.11.8) analysis. High quality reads were aligned to the *Fasciola hepatica* transcriptome (PRJEB25283) using HISAT2 (v.2.1.0) and transcripts generated from genome annotation (PRJEB25283) and counted using StringTie (v.1.3.6). Raw gene counts were reviewed for genes of interest (GOI) and analysed for differential expression using R version 3.6.2 and DESeq2 (v.1.26.0) package (FDR <0.001). A core annotation pipeline was developed to annotate GOI’s using BLASTx against NCBI non redundant protein (nr) database; predicted proteins generated using Transdecoder; BLASTp predicted proteins against uniprot reviewed sequence database; functional domain analysis interrogating PFAM and interproscan databases. Graphs were visualised using ggplot2, gate, pathview and upsetR R packages using custom scripts. Chart generated online at: https://app.lucidchart.com. Abbreviations; ivt=*in vitro*, ivv=*in vivo*.

Differentially expressed genes within these datasets offer insight into the biological differences of juveniles under distinct maintenance conditions and have the potential to identify ways in which *in vitro* culture methods could be improved. Since a primary phenotypic difference between *in vitro* and *in vivo* maintained juveniles is worm size, with age-matched *in vivo* juveniles being ∼15-times larger (Fig 2A-C), it can be hypothesized that differentially expressed genes play roles in the growth and development of juveniles; these genes have the potential to seed the identification of targets critical to pathogen virulence and establishment within the host. Staining proliferating cells of juveniles in both growth conditions using 5-ethynyl-2-deoxyuridine (EdU) revealed increased EdU+ cells in *in vivo* maintained juveniles (18859±1388 Edu+ cells after 18 h; n=4; Fig 2B) when compared to *in vitro* counterparts (316±30 Edu+ cells after 18 h; n=26; Fig 2A), revealing a 60-fold difference in the number of proliferating cells. The enhanced cell proliferation in *in vivo* juveniles was also evident when data were normalised to account for the ∼15-fold size difference (Fig 2C), confirming that the larger worms were supported by a ∼4-fold higher rate of cell proliferation.

**Fig 2:**
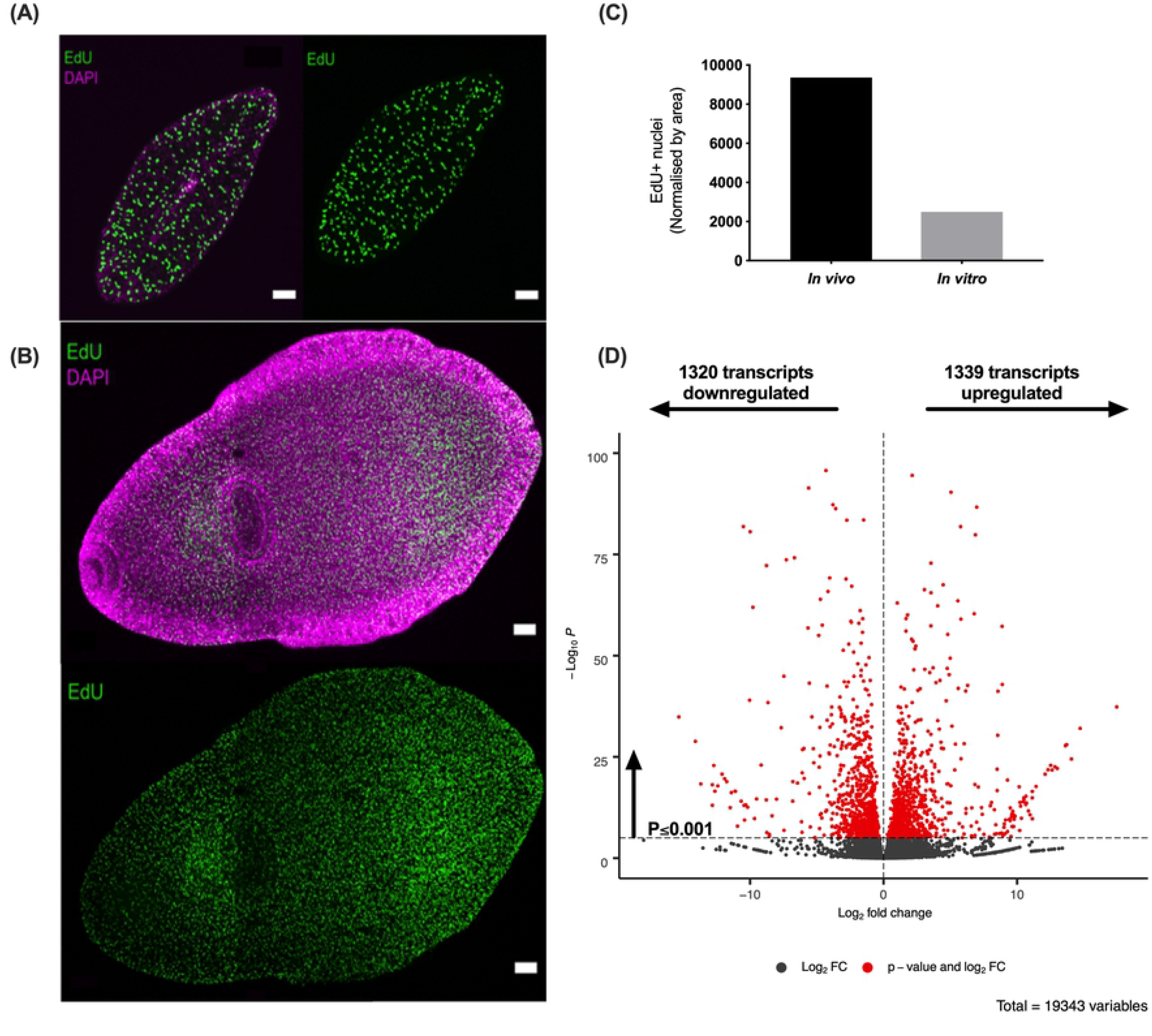
*In vitro* and *in vivo* maintained juvenile liver fluke display differences in growth and cell proliferation rate. **(A) Cell proliferation represented by EdU+ cells (green) of 21 day old (+1 day staining) *in vitro* maintained juvenile *Fasciola hepatica*. (B) Cell proliferation represented by EdU+ cells (green) of 21 day old (+1 day staining) *in vivo* maintained juvenile *F. hepatica*.** Confocal Z stack images; green= EdU+; magenta= DAPI; *in vitro* scale bar= 50 µm; *in vivo* scale bar= 100 µm. **(C) Proliferative cell count (EdU+) per mm^2^ between *in vitro* and *in vivo* maintained juveniles.** A ∼4-fold higher cell proliferation rate was seen in *in vivo* maintained juveniles. **(D) Volcano plot of differentially expressed transcripts between *in vitro* and *in vivo* maintained juveniles**. Differential expression analysis and plot generated using R version 3.6.2 and DESeq2 (v.1.26.0) package. Analysis identified 1339 transcripts upregulated and 1320 transcripts downregulated *in vivo* compared to *in vitro* maintained juveniles with a false discovery rate (FDR) of P≤0.001. Red = significantly differentially expressed transcripts. Grey = not differentially expressed transcripts.

### Differential expression of genes associated with selected cellular processes

DESeq2 was used to analyse differential expression of genes between *in vitro* and *in vivo* treatments. Analysis identified 1339 genes upregulated (6.9%) and 1320 genes downregulated (6.8%) *in vivo* compared to *in vitro* counterparts, amounting to a total of 13.7% of expressed genes being differentially expressed between maintenance treatments (Fig 2D). Annotations were assigned to 72% (1923) of differentially expressed genes from at least one hit match from interrogated databases of the annotation pipeline (S3 Table).

Notably, Gene Ontology (GO) term analysis highlighted an increase in cellular processes associated with DNA replication in datasets of *in vivo* maintained worms with an increased number of the genes associated with transcription regulation, DNA replication and microtubule based processes/movement compared to *in vitro* maintained worms (Fig 3A&B). Genes associated with nuclear based cellular components (nucleus, microtubules, MCM complex, chromosome, nucleosome) were also upregulated *in vivo* (Fig 3C). Although KEGG pathway analysis showed similar results, it more specifically highlighted a significant upregulation of genes associated with cell cycle, meiosis and cellular senescence pathways *in vivo* (Fig 3D) - all processes associated with cell division. GO term analysis of *in vitro* maintained juvenile datasets highlighted a more diverse range of gene functions. Genes associated with oxidation-reduction processes were upregulated in *in vitro* juvenile datasets suggesting some metabolomic differences between treatment groups, whilst an increased number of genes associated with transmembrane transport, G protein signalling and ion transport suggested an enhancement of some cell signalling pathways in *in vitro* juveniles when compared to *in vivo* counterparts (Fig 3A-C). KEGG enrichment analyses showed a significant downregulation of genes associated with the synaptic vesicle cycle, axon guidance and, more specifically, cholinergic signalling *in vivo* (Fig 3D). N glycan biosynthesis and protein processing in the endoplasmic reticulum were also significantly downregulated *in vivo* compared to *in vitro* maintained juveniles, interesting as glycosylation has been proposed to play a significant role in host-parasite interaction and suggests that glycosylation is an important protein modification within *in vitro* juveniles (Fig 3D). An increase in lysosomal pathway genes in *in vitro* maintained juveniles may be associated with the prevalence of cathepsin B proteases with greater homology to human proteases than the cathepsin L proteases that were more abundant in *in vivo* datasets. Cathepsin profiles are discussed in more detail later.

**Fig 3:**
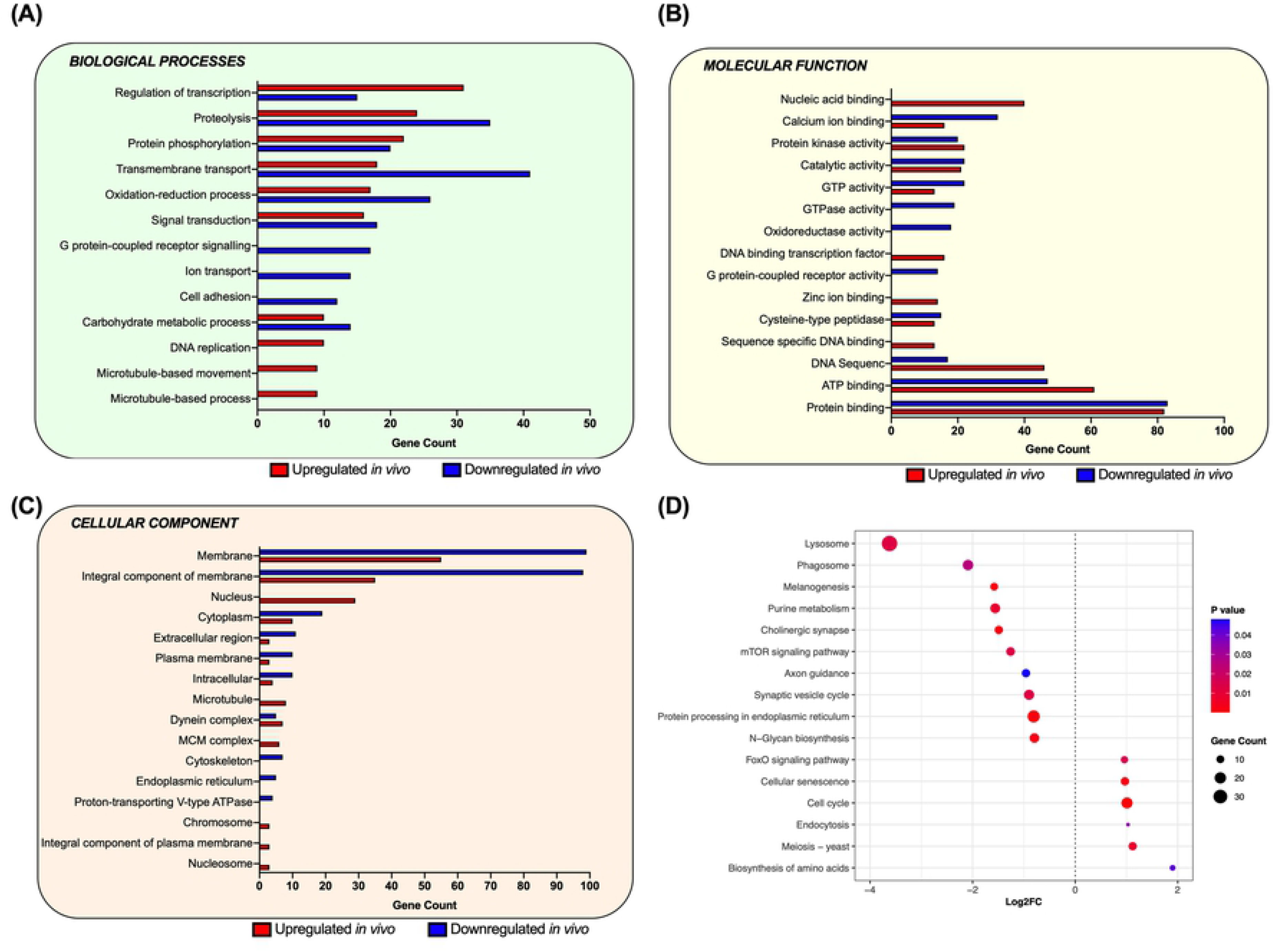
*In vivo* juvenile fluke display enhanced expression of genes associated with the cell cycle. **(A-C) Gene ontology (GO) term analysis of differentially expressed genes.** Gene counts associated with GO terms of (A) biological processes, (B) molecular function and (C) cellular components from differentially expressed and annotated genes on WormBase ParaSite (v.14). Upregulated datasets contain more genes associated with GO terms of cell cycle processes and DNA replication whilst downregulated datasets contain a greater number of genes associated with cell signalling and transport GO terms. Red represents the number of differentially expressed genes upregulated *in vivo*. Blue represents the number of differentially expressed genes downregulated *in vivo*. **(D) KEGG pathway analysis of differentially expressed genes.** KEGG pathways considered significantly differentially expressed as determined by statistical analysis using R (v.3.6.2), gage (v.2.36.0) and pathview (v.1.26.0) packages (P≤0.05). Dotted line = 0 log_2_FC; dots to the right of dotted line represent pathways upregulated *in vivo* and dots to the left of the dotted line represent pathways downregulated *in vivo*. Colour = *p* value; size = gene count.

### Cell cycle- and neoblast-associated genes are upregulated in in vivo maintained juveniles

Eighty nine percent (17,166) of total genes identified are found in all datasets (Fig 4A; 3x *in vitro=* 21d_ivt1-3; 3x *in vivo=* 21d_ivv1-3). In addition to differentially expressed genes, those genes which are present in only one treatment group, *i.e.* genes considered switched ‘on’ or ‘off’ *in vivo* are also genes of interest for understanding mechanisms of juvenile growth and development. Unsurprisingly, more genes (226; S3 Table) are present only in *in vivo* datasets, likely due to the demands of a more complex and changing host environment (S3 Table). 76/226 genes returned no hits when interrogated against the databases of the annotation pipeline, suggesting these genes are novel and unique to *in vivo F. hepatica* juveniles. ‘Hypothetical protein’ annotations were assigned to 65 genes and 85 genes were allocated function using the annotation pipeline described. A significant proportion of the genes only identified *in vivo* are associated with cell cycle processes and transcription regulation including cyclin-dependent kinase, cyclin, centromere protein, nucleosome assembly protein, transcription factors and zinc finger proteins (S3 Table). Also present only in *in vivo* datasets are genes associated with cell structure such as centrin, tubulins (alpha and beta chains) and actin associated proteins (slingshot protein phosphatase and actin-interacting protein) involved in microtubule and microfilament formation. Increased expression of these genes in the *in vivo* datasets is consistent with increased cell proliferation and turnover in these juveniles. Only 64 genes were present solely in *in vitro* datasets; 24 of these were assigned function using the annotation pipeline (S3 Table). Although no clear pattern of gene groups was evident, a histone 2B gene commonly associated with cell proliferation was within the *in vitro*-only datasets, suggesting that it could be associated with an earlier stage of growth and development in *F. hepatica* juveniles or is linked to an unknown feature of *in vitro* juvenile biology.

**Fig 4:**
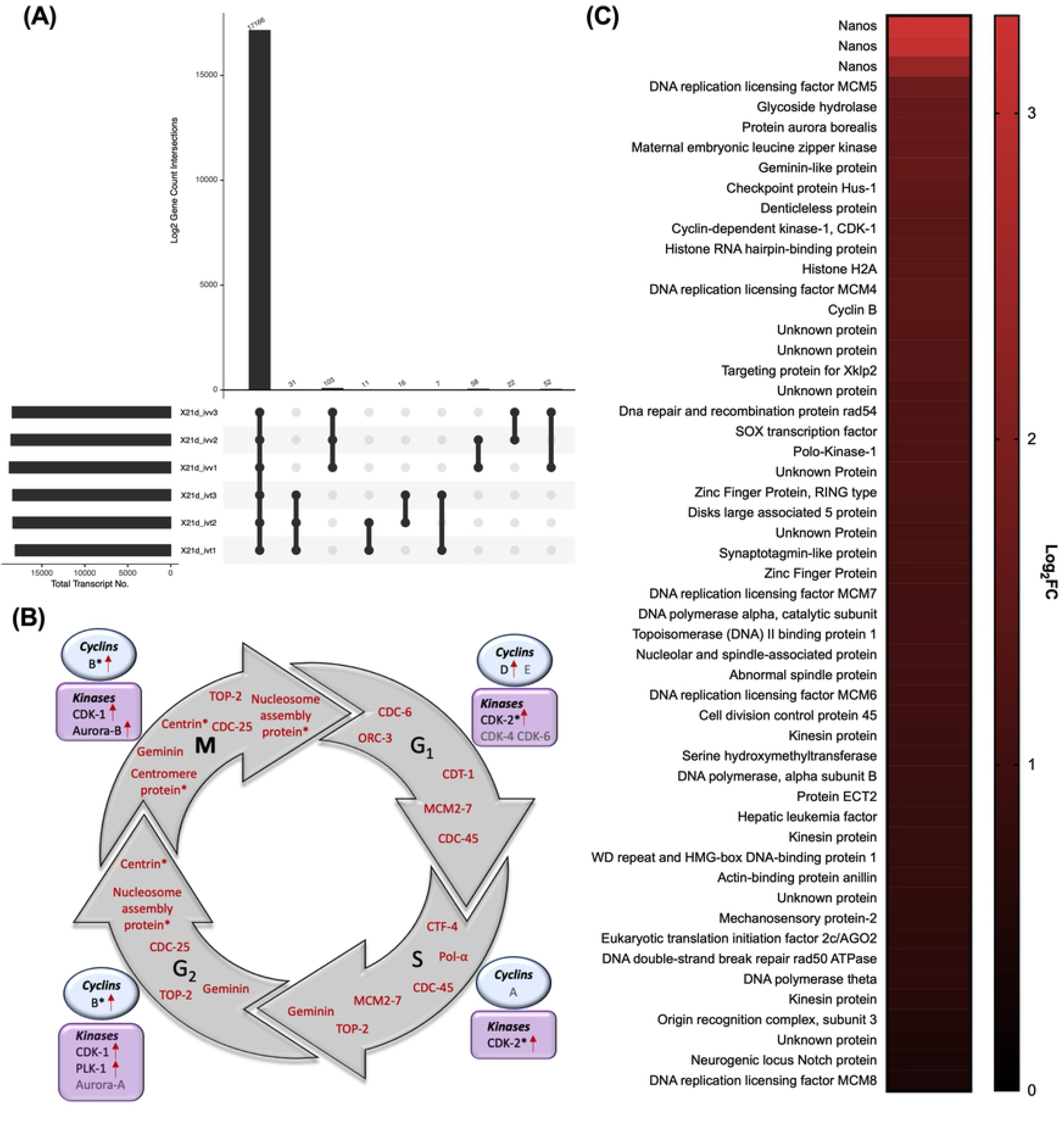
*In vivo* juvenile liver fluke displayed upregulation in cell cycle associated genes and neoblast-like stem cell markers. **(A) Upset plot showing spread of all transcripts across transcriptome replicates.** 19,343 protein coding genes were generated from transcriptome analysis. 17,166 genes were found in all generated transcriptomes. Genes of interest (GOI) include 65 genes present only in *in vitro* transcriptomes and 235 genes present only in *in vivo* datasets. Limit of analysis restricted to those genes present in at least 2 replicates of either transcriptome (i.e. *in vitro* or *in vivo*). **(B) Cell cycle associated genes upregulated *in vivo.*** Schematic showing phases and progression of the cell cycle (intermediate (G1), synthesis (S), growth (G2) and mitotic (M) phases). Important genes required for each cell phase to function in the human cell cycle and which have homologues in *Fasciola hepatica* that are upregulated *in vivo* are highlighted in red within the cycle. Those genes only present in *in vivo* datasets are marked with an asterisk (*). Key cyclins (in blue circles) and kinases (in purple squares) required for each phase of the cell cycle to progress in the human cell cycle are highlighted. *F. hepatica* homologues which are upregulated *in vivo* are marked with a red arrow. Important genes required for the cell cycle to function and progress are upregulated *in vivo*. Abbreviations: CDC(−6, −25, −45), cell division cycle; ORC-3, origin recognition complex subunit 3; CDT-1, chromatin licensing and DNA replication factor 1; MCM2-7, mini-chromosome maintenance complex; TOP-2, DNA topoisomerase 2; Pol-α, DNA polymerase alpha; CTF-4, chromosome transmission fidelity 4; CDK(−1, −2, −4, −6), cyclin-dependent kinase; PLK-1, polo-kinase 1. **(C) Neoblast markers upregulated *in vivo.*** Heatmap showing differentially expressed (log_2_FC) neoblast markers in *F. hepatica.* Neoblast markers identified as homologues to markers described by Collins *et al.* [24] in *Schistosoma mansoni*. 53/108 neoblast markers identified in *F. hepatica* were upregulated in *in vivo* datasets.

Closer examination of all cell cycle-associated genes shows a high proportion of these genes upregulated *in vivo* across all cell-cycle stages, corroborating an increased rate of cell division in these juveniles (Fig 4B). In particular, all components of the highly conserved MCM2-7 complex and cell division cycle genes (*cdc-6, cdc-45, cdc-25*) known to tightly regulate key components of DNA replication (Fig 4B). Key regulators of cell cycle progression, including cyclins and cyclin-dependent kinases (CDKs), are also significantly upregulated suggesting cell cycle activities are occurring at an increased rate *in vivo* (Fig 4B). The kinases polo-like kinase-1 (*plk-1*) and aurora-b kinase (*aurkb*), thought to ensure accurate chromosomal segregation, were also upregulated *in vivo*, which would appear to be consistent with the phenotypic observations (Fig 4B). To characterise further the relationship between growth/development and neoblast-like stem cells, differential expression analysis considered the 128 neoblast-like cell markers identified in *S. mansoni* [24]. Of these genes, 108 were found to have orthologues in the *F. hepatica* genome and 53 were shown to be differentially expressed and upregulated in *in vivo* maintained juveniles (S4 Table), consistent with the altered stem cell dynamics needed to drive faster growth and development in these juveniles (Fig 4C; S4 Table). Amongst the known neoblast markers significantly upregulated in the *in vivo* juveniles were the key transcriptional regulators *nanos* (3 genes) and *sox-1,* previously identified as being essential for neoblast proliferation in flatworms [24, 25] (Fig 4C). These data offer new insight into important genes regulating essential mechanisms of *F. hepatica* growth and development and provide a starting point for functional validation as potential control targets.

### Protease profile dynamics differ within the in vitro and in vivo maintained juveniles

Cathepsin (CAT) proteases represent >80% of the proteins secreted by adult liver fluke [26] and are of particular interest due to their role at the host parasite interface and potential in vaccine development [26–29]. CATs display marked temporal changes in expression during juvenile fluke development associated with altering proteolytic requirements of the parasite as it migrates within the mammalian host [26, 28]. 34 potential CATs (23 CATL and 11 CATB) have been previously described within the *F. hepatica* genome [26]. Our datasets, suggest that *F. hepatica* in fact express 43 individual CAT (29 CATL and 14 CATB) proteins (S4 Table). It should be noted that two novel gene sequences (MSTRG.20621; MSTRG.9158) identified through this analysis align well with previously annotated CATs (maker-scaffold10x_895_pilon-snap-gene-0.1; maker-scaffold10x_250_pilon-snap-gene-0.13), however, these novel sequences are significantly truncated at their N terminus suggesting they may have distinct functions. Annotated separately in this analysis, it is also possible these genes are isoforms of previously annotated genes which would bring the overall cohort of *F. hepatica* CAT genes down to 41 (27 CATL and 14 CATB). Of the 43 CATs identified from this analysis, 32 genes were differentially expressed suggesting that CAT profiles of *in vitro* and *in vivo* maintained juveniles are considerably different (Fig 5A). 22 CATLs were differentially expressed, of which 14 were upregulated *in vivo* (Fig 5A; left; S4 Table). One novel gene, MSTRG.9158 was identified as the most upregulated *in vivo* (Log_2_FC=17.46) and found to be a truncated CATL1 protein with high similarity to maker-scaffold10x_250_pilon-snap-gene-0.13. There was a general trend of CATL1 genes (including MSTRG.9158) being upregulated *in vivo*. SignalP (v.5.0) analysis of upregulated cathepsins showed 80% contained a signal peptide, suggesting they may be secreted at the host-parasite interface (S4 Table). CATL encoding genes downregulated *in vivo* are thought to belong to clade 5, suggesting these cathepsins in particular are more important in biology associated with *in vitro* maintenance (S4 Table). Ten CATB proteins were differentially expressed, 9 of these were downregulated *in vivo* 21 day-old worms (Fig 5A; left; S4 Table); 2 novel CATB proteins not previously annotated were identified (MSTRG.21893 and MSTRG.6836). The downregulation of CATB genes *in vivo* was also correlated with a downregulation of legumain regulatory peptidases, likely relating to the *trans*-activation of CATB proteases by legumains (Fig 5B). These data suggest CATB proteases are more important to the biology of *in vitro* maintained juveniles, whilst CATL proteases are more significant in the biology of *in vivo* maintained juveniles, likely due to host interactions. It is possible that the differences in cathepsin expression relate to developmental differences between the two treatment groups – indeed, a switch from CATB protease expression to CATL expression has been linked with the development of *F. hepatica* juveniles *in vivo* [26, 28]. To investigate this hypothesis the expression of CAT genes annotated in the genome (PRJEB25283, WBPS14; 21 CATL and 8 CATB genes) was correlated with expression in the previously published life stage transcriptomes, with a specific focus on newly excysted juvenile (NEJ), 24 hour juvenile and adult datasets (Fig 5A). These data show a general trend of CATs upregulated *in vivo* showing greater expression in the juvenile and adult stages of development, whilst the CAT genes downregulated *in vivo* show greater expression in the 24 hour NEJ stage of development (Fig 5A). These observations are consistent with a developmental delay in the *in vitro* maintained juveniles compared to their age matched *in vivo* juveniles. Anomalies in the pattern show some genes downregulated *in vivo* have higher expression in juvenile/adult datasets (CAT-B; maker-scaffold10x_1382_pilon-snap-gene-0.52 and maker-scaffold10x_889_pilon-snap-gene-0.31, CAT-L; maker-scaffold10x_231_pilon-augustus-gene-0.4) and some genes upregulated *in vivo* have greater expression in NEJ datasets (CATL; maker-scaffold10x_1853_pilon-snap-gene-0.17). This is likely due to the *in vitro* juveniles being significantly more developed than 24 hour NEJs and the *in vivo* juveniles expressing a mix of cathepsin B and cathepsin L proteins as they transition towards the adult life-stage. Other protease expression dynamics included an increased proportion of metalloproteinases in the *in vivo* juveniles, likely to aid in host tissue degradation (Fig 5B). There was an increased proportion of calpain proteases in the *in vitro* juveniles, consistent with observations in *Schistosoma* spp. where they are important for early juvenile migration and host evasion due to their role in immune defence [30].

**Fig 5:**
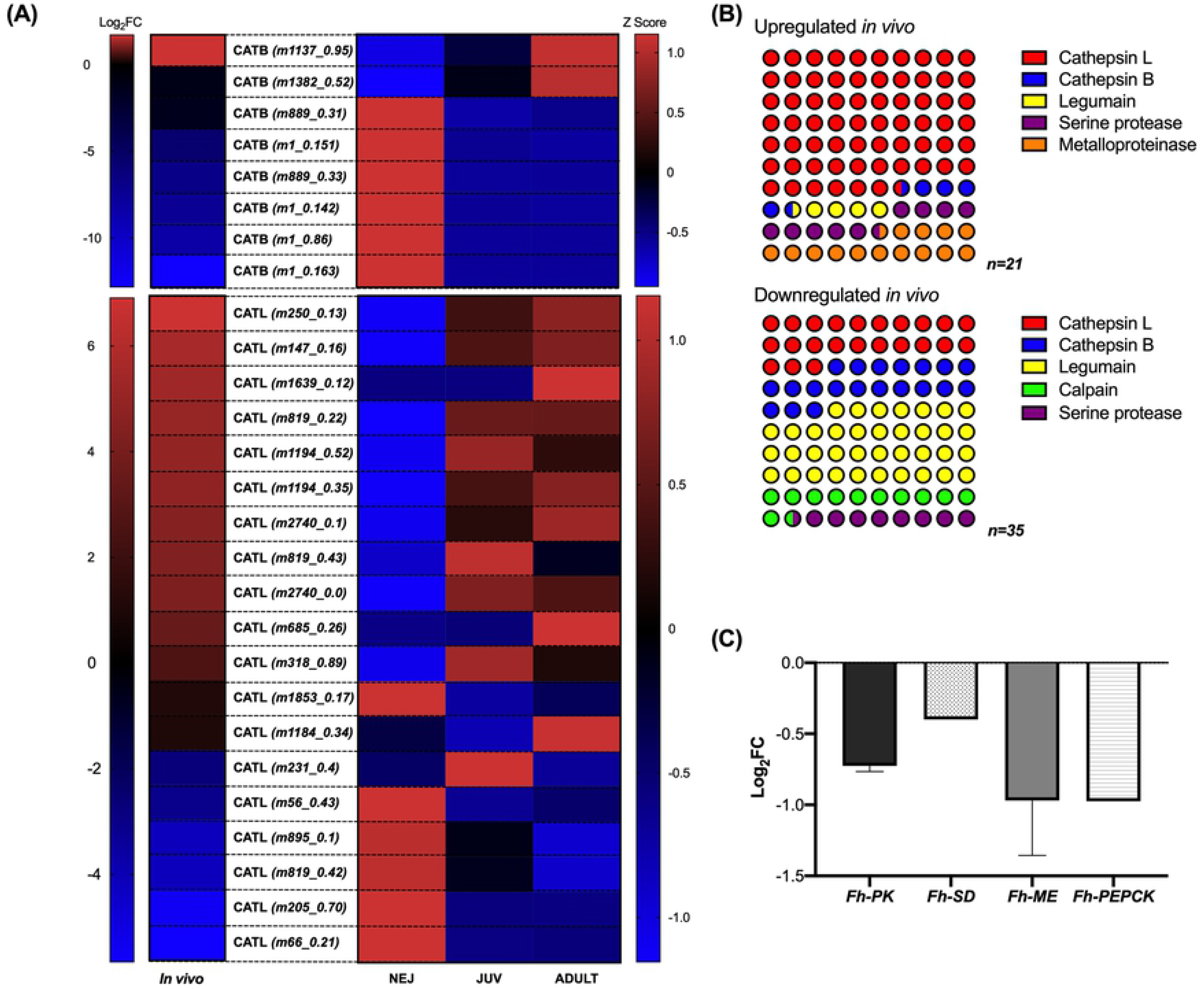
*In vivo* and *in vitro* juvenile liver fluke show divergent expression of proteases and metabolic enzymes. **(A) Cathepsin expression profile across newly excysted juvenile (NEJ), 21 day old juvenile (JUV) and adult *Fasciola hepatica*life stages compared to expression in *in vitro* and *in vivo* maintained *F. hepatica* juveniles.** Heatmap showing differential expression of cathepsin B (CATB) and cathepsin L (CATL) encoding transcripts in *in vivo* and *in vitro* maintained *F. hepatica* juveniles (log_2_FC) and across *F. hepatica* life stage transcriptomes generated by Cwiklinski *et al.* [20] (NEJ, JUV, ADULT; Z score of fragments per kilobase of transcript per million mapped reads, FPKM). Abbreviated gene IDs in brackets refer to genes in S5. Data show clear pattern of upregulation *in vivo* and greater expression in juvenile and adult transcriptomes, whilst downregulated genes *in vivo* show greater expression in NEJ transcriptome datasets. **(B) Protease/peptidase profiles of *in vitro* and *in vivo* maintained *F. hepatica* juveniles.** Proportion diagram shows greater expression of cathepsin L and metalloproteases in *in vivo* maintained *F. hepatica* juveniles whilst cathepsin B, legumain and calpain proteases show greater expression in *in vitro* maintained *F. hepatica* juveniles. **(C) Downregulation of key enzymes of aerobic and anaerobic carbohydrate metabolism in *in vivo* maintained *F. hepatica* juveniles.** Expression (log2FC) of key enzymes associated with metabolism in *in vivo* maintained *F. hepatica* juveniles. *Fh*-PK=pyruvate kinase and Fh-SD=succinate dehydrogenase are key enzymes of aerobic metabolism via Kreb’s cycle; *Fh*-ME-malic enzyme is a key enzyme of aerobic acetate production; *Fh*-PEPCK=phosphoenolpyruvatecarboxykinase, key enzyme of malate dismutation pathway.

### Metabolomic differences between in vitro and in vivo maintained F. hepatica juveniles

Parasitic flatworms rely heavily on carbohydrate substrates for energy metabolism [31]. *F. hepatica* have been shown to switch from aerobic energy metabolism via aerobic acetate production using the Tricarboxylic Acid Cycle (TCA) pathway, to anaerobic dismutation [20, 31] as they transition from free living to parasitic life stages. *In vitro* cultured juveniles are maintained under anaerobic conditions long term (5% CO_2_), but given the opportunity can also undergo aerobic metabolism, whilst 21 day *F. hepatica* juveniles burrowing through the liver parenchyma are undergoing a switch to predominantly anaerobic metabolism. Surprisingly, the key enzymes (pyruvate kinase, Fh-PK; succinate dehydrogenase, Fh-SD; malic enzyme, Fh-ME; phosphoenolpyruvate carboxykinase, Fh-PEPCK) associated with all pathways of carbohydrate metabolism (aerobic and anaerobic) are downregulated *in vivo,* suggesting enhanced metabolic activities, both aerobic and anaerobic, in the *in vitro* maintained juveniles (Fig 5C). The upregulation of lactate dehydrogenases in *in vitro* datasets suggests these juveniles are generating greater levels of lactate metabolic waste than *in vivo* juveniles, whilst the upregulation of acetate:succinate CoA-transferase in *in vivo* juveniles suggests that they are producing higher levels of acetate (S4 Table). To investigate this further, *in vitro* maintained juveniles were maintained in an anaerobic chamber, removing the ability to undergo aerobic respiration. These juveniles were unable to grow and showed declining health across a two week period when compared to juveniles maintained under 5% CO_2_ conditions (S5 Fig). After 3 weeks, significant death (87.4%) resulted in termination of the experiment.

### N-glycan biosynthesis and processing enzymes downregulated in vivo

KEGG pathway analysis highlighted the downregulation of genes associated with N-glycan biosynthesis and processing *in vivo* (Fig 3D). On closer examination of protein glycosylating genes described by McVeigh et al. [32], downregulation *in vivo* is particularly associated with components of the oligosaccharyl transferase (OST) complex involved in the *en-bloc* transfer of glycans to proteins [33], including catalytic core domain proteins STT3A/STT3B and RPN-1/-2 proteins (S5 Fig). Glycans are commonly linked to roles in host parasite interactions [34, 35] and the high expression of glycosylating enzymes associated with glycan biosynthesis in *in vitro* maintained worms suggests that aspects of host-parasite interactions are sustained *in vitro* even in the absence of a host. It is important to note that there was no differential expression of O-linked glycosylation genes, again thought to correlate with host-parasite interactions [34, 35].

### Components of the F. hepatica nervous system are downregulated in vivo

KEGG pathway analysis highlighted a significant downregulation of nervous system components *in vivo*. Flatworms employ both classical and neuropeptidergic neurotransmitters for neurotransmission [36–38] and both displayed markedly reduced expression in the *in vivo* juveniles (Fig 6).

**Fig 6:**
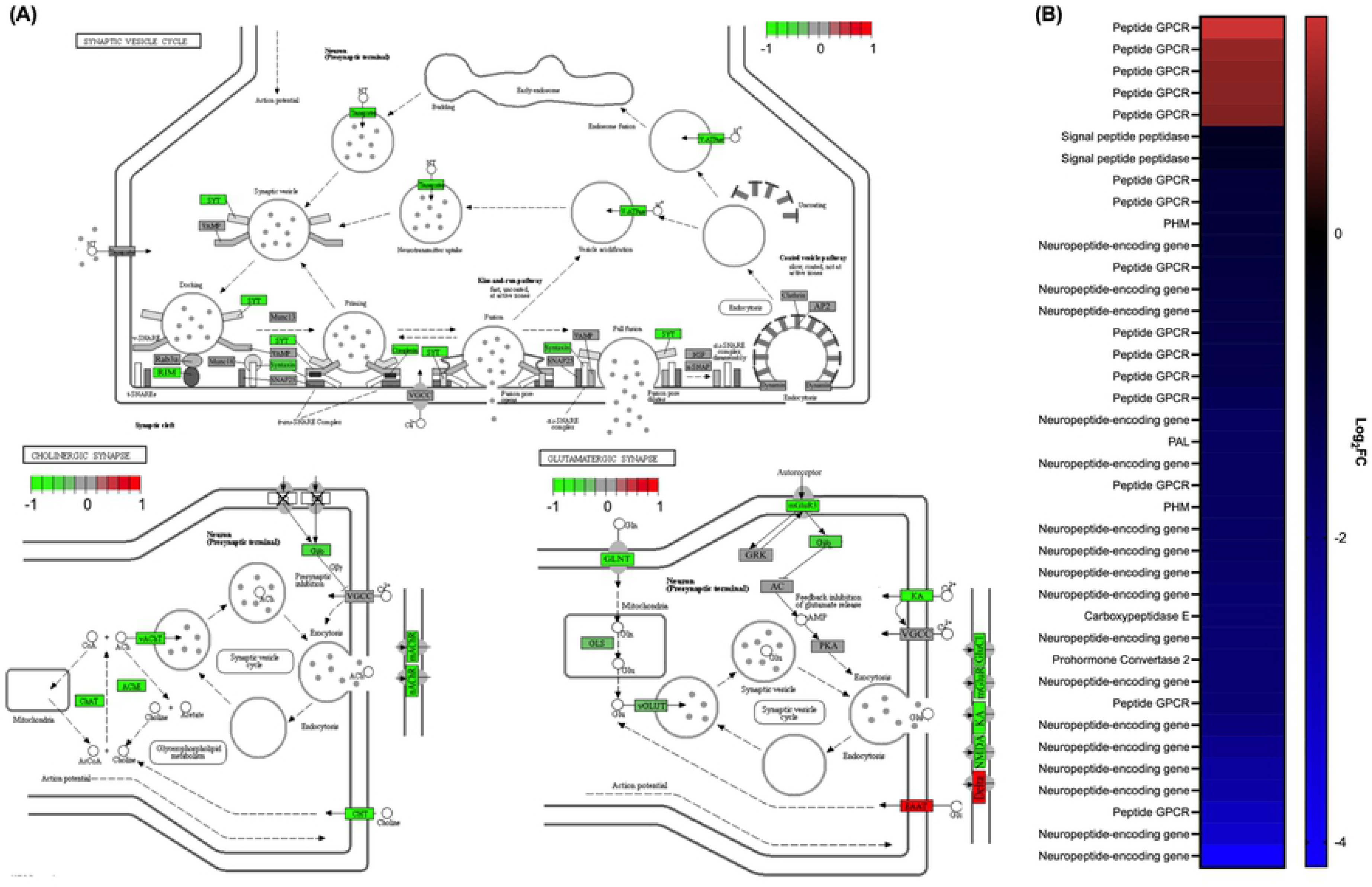
Key neuronal signalling pathways are downregulated in fast growing *in vivo* juvenile liver fluke. **Components of classical neurotransmission are downregulated in *in vivo* maintained juvenile liver fluke (A).** Analysis and diagrams generated using R (v.3.6.2), gage (v.2.36.0) and pathview (v.1.26.0) packages showing differential expression of nervous system components of the synaptic vesicle cycle, cholinergic signalling and glutamatergic signalling. *Fasciola hepatica* homologues of human components enclosed in rectangular boxes. Green=downregulated *in vivo,* red=upregulated *in vivo*, grey=not differentially expressed, X=not present in *F. hepatica* genome (PRJEB25283). Components are well conserved between human and *F. hepatica*species. KEGG pathway analysis identified a significant downregulation of the synaptic vesicle cycle and cholinergic signalling in *in vivo* maintained *F. hepatica* juveniles. Abbreviations: SYT, synaptotagmin; VAMP, vesicle associated membrane protein; Rab3a, ras-related protein; RIM, regulating synaptic membrane exocytosis protein; MUNC(−13, −18), syntaxin binding protein; SNAP25, synaptosome associated protein; VGCC, voltage-gated calcium channel; NSF, N-ethylmaleimide sensitive factor; αSNAP, NSF associated protein; AP2, adaptor related protein complex 2; ChAT, choline O-acetyltransferase; AChE, acetylcholinesterase; vAChT, vesicular acetylcholine transporter; CHT, high affinity choline transporter; Gi/o, guanine nucleotide-binding protein; mAChR, muscarinic acetylcholine receptor; nAChR, micotinic acetylcholine receptor; GLNT, amino acid transporter; GLS, glutaminase; vGLUT, vesicular glutamate transporter; GRK, G protein-coupled receptor kinase; AC, adenylate cyclase 1; PKA, protein kinase A; EAAT, glial high affinity glutamate transporter; mGluR, muscarinic glutamate receptor; GluCl, glutamate-gated chloride channel; KA, kainate type ionotropic glutamate receptor; NMDA, NMDA type ionotropic glutamate receptor; DELTA, delta type ionotropic glutamate receptor. **Components of neuropeptidergic signalling pathway are downregulated *in vivo* juvenile liver fluke (B).** Heatmap showing differentially expressed genes associated with neuropeptidergic signalling in *Fasciola hepatica.* All components of the neuropeptide processing pathway are downregulated *in vivo*. Our analysis identified 34 neuropeptide-encoding genes in the *Fasciola hepatica* genome (PRJEB25283) of which 17 are downregulated *in vivo* compared to *in vitro* counterparts. Our analysis further identified 44 peptide GPCRs, an additional 9 compared to the analysis carried out by McVeigh el al [44]. nine peptide GPCRs were downregulated *in vivo*, one of which is hypothesized to be an NPY receptor, whilst 4 were upregulated *in vivo*.

The synaptic vesicle cycle is responsible for packaging and releasing neurotransmitters from the synapse to modulate neurotransmission [39]. All major components of this pathway were significantly downregulated *in vivo* suggesting an increased rate of neurotransmitter release, and as a result, classical neurotransmission communication in *in vitro* maintained juveniles (Fig 6A; top). Further examination revealed downregulation of the cholinergic signalling pathway and a noteworthy partial downregulation of the glutamate-signalling pathway, emphasising the importance of these two pathways to the biology of *in vitro* maintained juveniles (Fig 6A; bottom). Although classical neurotransmission is most commonly associated with modulating neuromuscular control in parasitic flatworms [40], these data suggest a novel role in growth and development. The majority of genes associated with serotonin and dopamine classical neurotransmitter signalling pathways were not differentially expressed. Serotonin has been highlighted as an essential component of the *F. hepatica* nervous system with a core role in neuromuscular control [41, 42] - its stable expression across treatments suggests some core functioning of the liver fluke nervous system were not changed by maintenance treatment.

Interestingly, major components of neuropeptide signalling were also downregulated *in vivo* with the entire neuropeptide processing pathway displaying reduced expression [37]. Genes encoding two signal peptidases, prohormone convertase-2 (*PC-2*), carboxypeptidase E (*CPE*), peptidylglycine α-hydroxylating monooxygenase (*PHM*) and peptidyl-alpha-hydroxyglycine alpha-amidating lyase (*PAL*) were all downregulated *in vivo* (Fig 6B). The downregulation of neuropeptide processing suggests neuropeptides are being produced at a decreased rate *in vivo* compared with *in vitro* maintained juveniles, suggesting neuropeptide signalling displays changing expression dynamics during development. We identified 35 neuropeptide (*npp*)-encoding genes in the *F. hepatica* genome (PRJEB25283, WBPS14) and 17 *npp*-encoding genes were differentially expressed between maintenance treatments; all 17 were significantly downregulated in the *in vivo* juveniles compared to *in vitro* juveniles (S4 Table), corroborating the hypothesis that neuropeptide signalling is increased in the *in vitro* maintained juveniles and may play a previously undescribed role in modulating liver fluke growth and development. 6 *npp*-encoding genes were downregulated *in vivo* with a log_2_FC <-2 and are of particular interest as potential modulators of growth and development in *F. hepatica* juveniles. Two of the most significantly downregulated *npp*-encoding genes are NPF/NPY neuropeptides, homologous to vertebrate NPY neuropeptides (S4 Table) [38, 43]. McVeigh et al. [44] identified 35 *F. hepatica* G-protein coupled receptors (GPCRs). Combining available genomic and transcriptomic datasets suggests *F. hepatica* expresses 44 peptide GPCRs, of which 15 are differentially expressed (S4 Table). Ten peptide GPCRs were downregulated *in vivo* and one of these was previously hypothesized to be an NPY receptor. Further, 5 peptide GPCRs were downregulated in the *in vitro* juveniles, corroborating KEGG pathway analysis which suggests GPCR signalling is relatively higher in the *in vitro* maintained juveniles. These observations support the hypothesis that selected classical neurotransmitters and neuropeptides act to modulate growth and development of *F. hepatica* juveniles, warranting further experimental exploration.

### MicroRNAs (miRNA) are differentially expressed in in vitro and in vivo maintained juveniles

This study identified 103 miRNAs in liver fluke, including 14 novel miRNAs (S6 Table) and 89 miRNAs reported in previous studies on *F. hepatica* [45–49]. Recently, Herron et al [49] highlighted considerable redundancy across published miRNAs and moved to refine the current cohort of F*. hepatica* miRNAs by removing duplicate sequences of high similarity and retaining the longer sequence as individual miRNAs. Applying this approach to our dataset refines our final miRNA dataset to 89 miRNAs (75 previously published [45–49]; 14 novel). Of the 89 total miRNAs described for *F. hepatica*, 31 were differentially expressed between *in vivo* and *in vitro* maintained juveniles; 18 miRNAs were significantly upregulated in *in vivo* juveniles and 13 were significantly downregulated in *in vivo* juveniles, suggesting a role for these miRNAs in transcriptional regulation across maintenance conditions (Fig 7A). Gene ontology (GO) term analysis of predicted gene targets for differentially expressed miRNAs shows the increased expression of mRNA targets associated with transcriptional regulation, microtubule-based processes and DNA replication, whereas predicted mRNA targets associated with transmembrane transport, ion transport and signal transduction were downregulated in *in vivo* juveniles (Fig 7B). These data correlate with mRNA analysis and suggest these processes are at least partially regulated by miRNAs in liver fluke. Notably, Wnt signalling was identified as a process likely to be impacted by differentially expressed miRNAs with upregulated miRNAs (*fhe-mir-10-5p*, *fhe-mir-190-5p*, *fhe-mir-2b-3p*, *fhe-pubnovelmiR-23-3p*, *fhe-novelmir-50-5p*, *fhe-novelmir-28-3p*) predicted to target Wnt-associated genes that displayed reduced expression in *in vivo* maintained juveniles (Fig 7, S6 Table). Five Wnt genes downregulated in *in vivo* juveniles were predicted as regulated by miRNAs (2 Wnt proteins, maker-scaffold10x_735_pilon-snap-gene-0.38 & maker-scaffold10x_254_pilon-augustus-gene-0.12; 1 secreted frizzled-related protein, maker-scaffold10x_405_pilon-snap-gene-0.20; 2 frizzled GPCRs, maker-scaffold10x_541_pilon-snap-gene-0.38 & maker-scaffold10x_944_pilon-snap-gene-0.48). Three downregulated miRNAs (*fhe-novelmir-48-3p*, *fhe-mir-184-5p* & *fhe-pubnovelmir-22-3p*) were also predicted to target low-density lipoprotein receptor-related protein (LRP) 5/6 (maker-scaffold10x_442_pilon-snap-gene-0.18), a Wnt-associated gene upregulated in *in vivo* maintained juveniles (Fig. 7, S6 Table).

**Fig 7:**
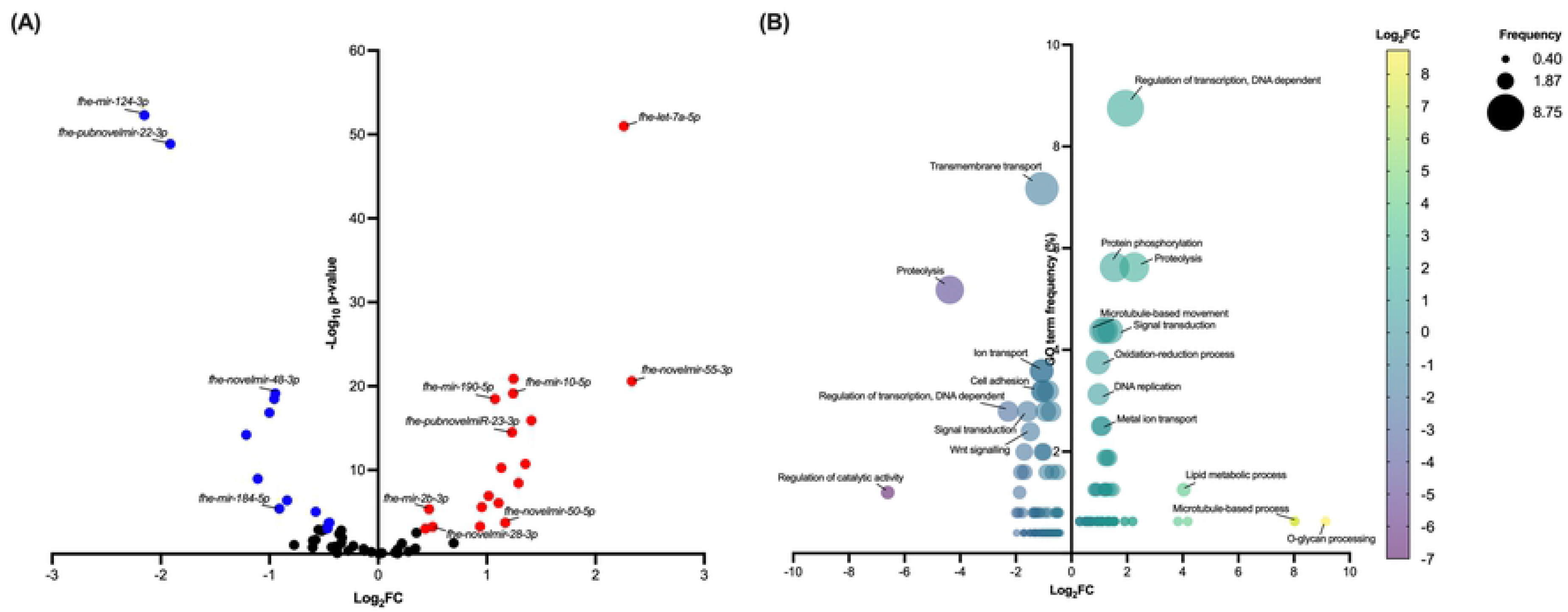
Differences in miRNA profiles of *in vitro* and *in vivo* juveniles support their role in liver fluke developmental processes. **(A) Differential expression of micro (mi)RNA sequences in *in vitro* and *in vivo* maintained 21-day old *Fasciola hepatica* juveniles.** Analysis identified 31 differentially expressed miRNAs between treatment groups (false discovery rate; P≤0.001). Red points highlight significantly upregulated miRNAs *in vivo*, whereas blue points highlight significantly downregulated miRNAs *in vivo*. The two most significantly upregulated miRNAs (*fhe-let-7a-5p,* first identified by Fromm et al. [46] and *fhe-novelmir-55-3p*), downregulated miRNAs (*fhe-mir-124-3p*, first identified by Fontenla et al. [47] and *fhe-pubnovelmir-22-3p* first identified by Fromm et al. [46]) and miRNAs identified as associated with Wnt signaling (*fhe-mir-10-5p*, *fhe-mir-184-5p*, *fhe-mir-190-5p*, *fhe-mir-2b-3p*, *fhe-pubnovelmir-22-3p*, *fhe-pubnovelmiR-23-3p*, *fhe-novelmir-48-3p*, *fhe-novelmir-50-5p* & *fhe-novelmir-28-3p*) are labelled. Analysis carried out using DESeq2 (v.1.26.0). **(B) Gene ontology (GO) terms associated with biological processes of differentially expressed miRNA predicted gene targets.** GO terms retrieved from WormBase ParaSite (WBPS15) for predicted miRNA targets upregulated or downregulated *in vivo*. Number of individual GO terms associated with biological processes (percentage frequency of total GO terms) plotted alongside average Log_2_ fold change of predicted transcripts from differential expression analysis of *in vitro* and *in vivo* maintained *F. hepatica* juveniles. Bubble colour = transcript expression (Log_2_FC); bubble size= GO term frequency (percentage gene counts).

Two downregulated miRNAs were more significantly differentially expressed than others, *fhe-mir-124-3p* identified by Fontenla et al. [47] and *fhe-pubnovelmir-22-3p* identified by Fromm et al. [46] (Fig 7A). In contrast *fhe-let-7a-5p,* first identified by Fromm et al. [46] and considered a highly conserved miRNA, was the most significantly upregulated miRNA in *in vivo* maintained juveniles, whilst *fhe-novelmir-55-3p* identified from this study was the most upregulated miRNA (Log_2_FC= 2.33) in the *in vivo* maintained juveniles (Fig 7A). Analysis suggested that these four miRNAs play a significant role in regulating the expression of genes associated with *in vitro* and *in vitro* survival. Differentially expressed predicted targets for these miRNAs include transcription factors, cell cycle proteins, apoptosis inhibitors, growth factors, metabolomic and glycosylating enzymes (S4 Table).

## Discussion

To date, this is the most in depth transcriptomic study of *F. hepatica* juveniles, providing crucial datasets for understanding the biology of this pathogenic stage, improving understanding and informing drug target identification and validation efforts.

13.7% of genes were considered differentially expressed in this study, with >86% of genes expressed at similar levels in *in vitro* and *in vivo* maintained juveniles. This suggests that core biological functioning is consistent in juveniles under the two growth conditions and despite the size differences observed between juveniles, the current *in vitro* maintenance platform supports relevant biological processes [15]. This corroborates previous observations that *in vitro* maintained juveniles develop phenotypic attributes consistent with *in vivo* developing juveniles [15], supporting the value of the *in vitro* functional genomics platform for initial drug target validation studies prior to the use of animal models of infection. Although *in vitro* and *in vivo* juvenile biology appear largely consistent, it is clear that parasites are highly adaptive and respond rapidly to changing external conditions [50]. It is likely that *in vitro* maintained juveniles are not exposed to host-related triggers that enhance growth/development dynamics as observed in the *in vivo* juveniles. Studies of nematode parasites suggests that parasites undergoing tissue migration are likely to be larger than closely related species that do not migrate and showcase greater levels of growth and development, suggesting the action of migration itself can stimulate this biology [51].

Transcriptome differences between the *in vitro* and *in vivo* juveniles were consistent with the observed differences in the rate of cell proliferation. Genes associated with cell proliferation and known neoblast markers were significantly upregulated *in vivo*, correlating with larger juvenile size. Neoblasts have previously been associated with growth and development of many flatworm species, including *F. hepatica* and related *Schistosoma mansoni* [15, 24, 52-55]. In the cestode *Echinococcus multilocularis*, stem cells were shown to be the only cells driving metacestode growth and regeneration [55]. Again, this suggests proliferation at a higher rate *in vivo* leading to more rapid development of these juveniles. The identification of cues that further stimulate cell proliferation is key to improving the current *in vitro* culture methods. In addition to the role in growth and development, neoblast proliferation has also been associated with rapid tegumental renewal and tissue repair of parasites *in vivo*, suggesting the importance of these cells to host-parasite interactions [56]. The tegument is the barrier between host and parasite such that its rapid renewal allows parasites to counter host immune responses through active evasion and/or repair of immune response-associated damage [56]. Clearly, the upregulation of neoblast-associated genes *in vivo* may partly relate to the demands of immune challenge and *in vivo* survival, challenges not faced during *in vitro* culture. It is noteworthy that a higher rate of neoblast proliferation was associated with the mitigation of tissue damage and crucial to *S. mansoni* survival *in vivo* [57].

Metabolomic differences in *in vitro* and *in vivo* juveniles may also help explain the observed differences in juvenile growth and development. Surprisingly, highly expressed genes associated with carbohydrate metabolism (aerobic and anaerobic) were upregulated in *in vitro* maintained juveniles. It is possible that this observation relates to the fact that chicken serum has a glucose content twice the level of that seen in mammals [58]; the increased expression of genes associated with carbohydrate metabolism may reflect the juveniles taking advantage of the higher substrate availability. During their development from NEJs to adult fluke, juvenile *F. hepatica* are reported to display a shift from aerobic to anaerobic carbohydrate metabolism. Early studies by Tielans *et al*. showed under aerobic conditions that juveniles transitioned from metabolism dominated by the Krebs cycle in early parenchymal stages, to aerobic acetate production dominating during later parenchymal stages, with malate dismutation being dominant in the bile duct stages [59]. Although specific to aerobic conditions and likely not a true representation of *in vivo* scenarios, our data appear consistent with a similar pattern of development such that *in vitro* juveniles are undergoing metabolism more akin to earlier parenchymal stages utilising glycolysis to produce high levels of lactate, whilst the *in vivo* juveniles are generating acetate, akin to later parenchymal stages. Note that culturing juveniles under anaerobic conditions led to the death of juveniles suggesting that even *in vivo,* migrating juveniles continue aerobic metabolism. These observations were supported by the fact there were no distinct differences in the expression of aerobic and anaerobic metabolism genes between treatment groups.

The slower development of *in vitro* juveniles is also reflected in the expression patterns of CATL and CATB proteases. It is well documented that juvenile fluke undergo a developmental shift from CATB expression to CATL expression as they develop from NEJs in the duodenum to adult liver fluke in the bile duct [26–28]. This is thought to reflect the changing protease requirements associated with feeding and tissue degradation [26]. Our transcriptomic datasets show greater expression of CATB proteases in the *in vitro* juveniles, reflective of early stage *F. hepatica* juveniles, whilst CATLs are more prominent in the *in vivo* juveniles, characteristic of later stages of development. The cathepsin profile developmental shift was previously described as occurring in the absence of host signalling [20], suggesting that the developmental delay of *in vitro* maintained *F. hepatica* juveniles is responsible for this difference. It is also possible that host-derived triggers during migration drive the changing cathepsin expression profiles seen in the *in vivo* juveniles. These data encourage further studies comparing earlier *in vivo* juveniles and later *in vitro* juveniles to see if matching cathepsin expression profiles could be a proxy for the alignment of *in vivo* and *in vitro* developmental stage. In addition to cathepsins, other proteases thought to play roles in host-parasite interactions, such as calpains, were also evident in the *in vitro* datasets [30]. Somewhat surprisingly, the proposed immunomodulatory protein helminth defence molecule (HDM-1), was not differentially expressed between the *in vitro* and *in vivo* maintained juveniles [60–62]. Indeed, the gene encoding HDM-1 had a raw gene count of >1 million in both treatment groups.

Glycan biology is also considered key for host-parasite interactions [34]. Our analysis revealed the consistent expression of proteins associated with O-linked glycosylation and the higher expression of N-linked processing enzymes in *in vitro* maintained juveniles. The specific relationship between N-glycans and *in vitro* maintained juveniles is unknown, but it is perhaps due to the availability of glucose in chicken serum, an important precursor of the N-linked glycan biosynthetic pathway [63]. It is also possible that these specific glycans are more important for some aspects of biology associated with *in vitro* culture of juveniles or are associated with host-parasite interactions at earlier stages of parasite development. These observations suggest that *in vitro* cultured *F. hepatica* juveniles have utility in informing aspects of the biology of genes involved in host-parasite interplay.

Major components of both classical and neuropeptidergic signalling pathways in the liver fluke nervous system were downregulated *in vivo,* including synaptic vesicle cycle components and all elements of the neuropeptide processing pathway, indicating that these pathways are more prominent in the biology of *in vitro* maintained worms. Acetylcholine signalling in particular was a significantly upregulated classical neurotransmitter pathway in *in vitro* maintained juveniles. Notably, both acetylcholine and NPF/NPY neuropeptides have been identified as functioning in behaviours associated with nutrient acquisition and feeding in invertebrate species [64–66]. Acetylcholine signalling has been shown to function in the modulation of glucose transport in *S. mansoni* [65], whilst Drosophila NPF functions downstream of insulin signalling to regulate larval feeding [67]. The upregulation of these pathways may reflect the contrasting nutrient availabilities, resulting in changing behaviours of nutrient acquisition that could impact the growth and development of juveniles.

Of particular interest is the hypothesis that these neuronal signalling systems play important roles in the modulation of growth and development of juvenile *F. hepatica* via mechanisms that influence stem cell dynamics and/or nutrient acquisition. Recent advances in cancer biology identifies acetylcholine receptors as key players in cancer development due to their modulatory role in cell proliferation and expression in non-neuronal cell types [68]. Receptors for acetylcholine have been identified in all human stem cell populations and in differentiated and undifferentiated cells types, supporting an important function beyond basic nervous system functioning towards determination of cell fate and proliferation [69, 70]. The role of acetylcholine signalling in the growth and development of invertebrate species is unclear, although as early as 1985, the exogenous application of acetylcholine was reported to improve the growth and proliferation of *Drosophila* cell lines *in vitro* [71]. The activation of nicotinic acetylcholine receptors by the nicotinic agonist dimethylphenylpiperazinium (DMPP) resulted in delayed cell division and differentiation, stunting the development of *Caenorhabditis elegans* L2s via interaction with the insulin/IGF and Ras-MAPK growth signalling pathways [72]. Further, *in situ* hybridisation experiments showed the extensive expression of nicotinic acetylcholine receptors in developing *Brugia malayi*, suggesting a specific role in embryogenesis and spermatogenesis [73]. NPF/NPY neuropeptides have also been linked to the modulation of regeneration and germline cell proliferation in invertebrates [74–76]. Non-neuronal NPF neuropeptides were shown to regulate bone morphogenetic protein (BMP) signalling and act as a key regulator of mating-induced germline cell proliferation in *Drosophila* females [76]. In free living flatworms, NPF was found to accelerate pharyngeal regeneration in *Girardia tigrina* [77], whilst RNAi of prohormone convertase 2 (PC2) and non-neuronal NPY-8 had significant impacts on the reproductive maturation of *Schmidtea mediterranea*, suggesting neuropeptides are essential for germ cell differentiation [78]. It is therefore possible that differential expression of these neuronal signalling pathways relates to a fundamental role of the nervous system in regulating the growth and development of *F. hepatica* juveniles; the data would support the hypothesis that cell proliferation and/or related growth mechanisms are inhibited by a cadre of neuronal signalling molecules. It is also possible that an upregulation of neuronal signalling in *in vitro* juveniles may result from their maintenance in an unfamiliar environment lacking directional cues, heightening the expression of systems involved in host/niche finding behaviours.

miRNAs are an abundant class of regulatory genes that control many cellular and developmental processes [79]. Analysis of the miRNA complements of *in vitro* and *in vivo* maintained juvenile *F. hepatica* supports a significant role for these small RNAs in the modulation of developmental processes, including transcription and translation. The most significantly upregulated miRNA in *in vivo* juveniles was *fhe-let-7a-5p;* this miRNA is highly conserved across diverse organisms and has a significant role in the modulation of stem cells and the promotion of cell differentiation [80, 81], consistent with its upregulation in the more highly developed *in vivo* maintained *F. hepatica* juveniles. *Fhe-mir-124-3p* plays a key role in neuronal cell differentiation [82] and is also thought to be a growth suppressor [83], potentially correlating with its higher expression in the smaller *in vitro F. hepatica* juveniles. The downregulation of *fhe-mir-124-3p* in *in vivo* juveniles may reflect the developmental stage of these juveniles with well-formed neuronal systems supported by comparatively lower levels of neuronal development. It was notable that GO term analysis of predicted miRNA targets largely correlated with the most significantly differentially expressed mechanisms identified through mRNA transcriptome analysis. In addition, Wnt signaling was identified as a process regulated by differentially expressed miRNAs leading to an overall downregulation of Wnt mRNAs *in vivo.* miRNAs are well established regulators of Wnt signaling, a process essential for early organism development [84]. Downregulation of specific Wnt proteins, secreted frizzled-related proteins and frizzled GPCRs *in vivo,* suggest these proteins in particular have key roles in early stage developmental processes, consistent with later stage juveniles maintaining higher levels of tissue differentiation. Upregulation of low-density lipoprotein receptor-related protein (LRP) 5/6, a Wnt-associated gene, in *in vivo* juveniles suggests this protein is involved in later stage developmental processes.

Overall, this study has provided unique insight into the biology of a helminth parasite maintained *in vitro* and encourages the exploitation of *in vitro* cultured parasites in functional genomics studies to inform aspects of *in vivo* biology and control target validation. Where biology differs in relation to growth and development, new insights into metabolomic differences offer a basis for improving *in vitro* culture methods towards identifying key triggers for adult fluke development. Most of the transcriptomic differences between *in vitro* and *in vivo* maintained juvenile fluke relate to their divergent growth rate/stem cell proliferation and the developmental differences seen in the two groups of parasites. The data expose a key role for miRNAs in coordinating the developmental differences seen in *in vivo* (fast growing) and *in vitro* (slow growing) juveniles. Further, the data highlight dramatic changes in the expression of neuronal signalling systems, consistent with a role for the nervous system in suppressing the growth and development of *F. hepatica* juveniles. The observations encourage the search for new potential control targets associated with signalling systems that regulate juvenile growth/development in liver fluke.

## Materials and Methods

### Ethical statement

This work was carried out in accordance with the Animals (Scientific Procedures) Act 1986 adopting the principles of the 3Rs (Replacement, Reduction and Refinement). The Animal Project Licence was: PPL 2764. The methods proposed under the licence were approved by the QUB Animal Welfare Ethical Review Body and animals were euthanised using carbon dioxide gas.

### Fasciola hepatica material

Italian strain *F. hepatica* metacercariae were purchased from Ridgeway Research for generation of transcriptomes. For *in vivo* maintained *F. hepatica*, 16 Sprague-Dawley rats were infected with 25 metacercariae each and maintained for 21 days before fluke were recovered from liver tissue. Livers were cubed and incubated at 37°C, 5% CO_2_ to allow juvenile fluke to emerge for collection. All juveniles were collected within 2 hours of initial liver processing, washed five times in RPMI to remove liver material and snap frozen for storage at −80°C prior to further processing. One biological replicate contained 9 juveniles from at least 3 rats. *In vitro* maintained juvenile replicates were manually excysted as described by McVeigh et al. with a prior preparation step of removing the outer casing of metacercariae and bleaching for 2-3 minutes [85]. Excystments of replicates were carried out on consecutive days to generate a total of 1000 juveniles per replicate. Juveniles were maintained in 50% chicken serum and RPMI as described by McCusker *et al.* for 21 days prior to being snap frozen for storage at −80°C prior to further processing [15]. Juveniles maintained under anaerobic conditions were placed in an anaerobic chamber (Don Whitley Scientific, Shipley, UK) set to 37°C and supplied with anaerobic mixed gas (10% CO2, 10% H2 in N2, BOC). To facilitate the maintenance of anaerobic conditions, media were incubated under anaerobic conditions for at least 4 h prior to media changes which were carried out daily. A time-matched control group was maintained as standard with 5% CO2. The pH of anaerobic media was checked to ensure that increased CO2 levels did not render it significantly more acidic than media incubated at 5% CO2. Trials ran for a duration of 3 weeks, with worm survival being monitored on a weekly basis. Worm death was defined by a total lack of movement and darkened appearance.

### Labelling proliferative nuclei with 5-ethynyl-2-deoxyuridine (EdU)

Visualisation of *Fasciola* proliferative cells from 21 day old juveniles maintained *in vitro* and *in vivo* was achieved by labelling nuclei undergoing DNA-synthesis with 5-ethynyl-2-deoxyuridine (EdU; ThermoFisher Scientific) as described by [15]. Incubations were carried out at a final concentration of 500 µM in 50% chicken serum for 24 hours. Juveniles were then flat fixed in 4% paraformaldehyde under coverslips for 10 minutes (*in vitro*) and 40 minutes (*in vivo*) followed by 4-hours free-fixing at room temperature. EdU incubated, fixed juveniles were processed for detection using the Click-iT EdU Alexa Fluor 488 imaging kit, as per kit instructions (ThermoFisher Scientific). Background labelling of all nuclear DNA was achieved using 4′,6-diamidino-2-phenylindole (DAPI). Samples for analysis were mounted in Vectasheild (Vector Laboratories) and viewed on Leica TCS SP5 or SP8 confocal microscopes as maximally projected z-stacks generated from 12-15 optical sections from ventral to dorsal surface.

### RNA extraction, library preparation and sequencing

3 biological replicates of *in vitro* maintained and *in vivo* retrieved juveniles were prepared. RNA was extracted using TRIzol reagent (Life Technologies) with an isopropanol precipitation, using glycogen as a carrier. Precipitation was performed overnight at −20°C to maximise small RNA recovery. RNA was DNase treated using the Turbo DNase kit (Ambion) following manufacturer’s instructions and quality control was performed using bioanalyzers to assess for degradation; Qubit for accurate quantification and NanoDrop to indicate sample purity. All accepted samples displayed a 260/280 >2, *i.e*. pure RNA with no protein contamination, and a 260/230 >1.8. Sample library preparation and sequencing were carried out by the Centre for Genomic Research at the University of Liverpool as follows. For RNAseq, libraries were prepared using a PolyA selection and the NEBNext Ultra Directional RNA library kit for Illumina to prepare dual indexed, strand specific libraries. Paired end sequencing (2×150bp) was performed on the 6 libraries (3 *in vivo*, 3 *in vitro*) using the Illumina HiSeq 4000 platform and generated in excess of 280M mappable reads (∼47M reads per sample). For small-RNAseq, libraries were prepared using the NEBNext Small RNA library preparation kit for Illumina and single-end sequencing (1×50bp) performed on the HiSeq2500 platform.

### Assembly, annotation and sequence data analyses

Raw sequences were trimmed for the presence of Illumina adapter sequences using Cutadapt (v.1.2.1) [86] and low quality reads using Sickle (v.1.200) [87] with a minimum window quality score of 20. Remaining reads under 10bp were removed. Final quality control was performed using FastQC (v.0.11.8) [88] with default parameters. The *F. hepatica* genome contigs (PRJEB25283) and associated GFF file (PRJEB25283) were downloaded from WormBase ParaSite (WBPS14; https://parasite.wormbase.org/index.html) [89] and annotations were converted to GTF format using Cufflink (v.2.2.2.20150701) [90]. Trimmed FastQT files were aligned to the *F hepatica* genome (PRJEB25283) using HISAT2 (v.2.1.0) [91] with default parameters and transcripts compiled and counted using StringTie (v.1.3.6) [92, 93]. Annotated transcripts were combined to generate a gene count file, removing transcript isoforms. Gene count datasets were filtered to remove non-coding genes and reads with zero gene counts. Data were normalised for sequencing depth and RNA composition using DESeq2 (v.1.26.0) [94] package in R (v.3.6.2) [95] with default parameters.

### Annotation and analysis of genes of interest

Raw gene count datasets generated as previously described were mined for genes absent in one treatment group (across 3 replicates of *in vitro* or *in vivo* transcriptomes) and present in at least 2 replicates of the opposite treatment group with a total gene count ≥10. These genes were designated as switched ‘on’ or ‘off’ *in vivo*. Differential expression analysis of protein coding genes was quantified using the DESeq2 (v.1.14.1) [84, 96] package in R (v.3.6.2) [95]. A p-value threshold was set for a false discovery rate (FDR) of <0.001 and no threshold was applied for fold change differences. Genes identified using these parameters were described as upregulated or downregulated *in vivo*. Differentially expressed genes were annotated as described in Fig 1. BLASTx analysis of gene sequences was carried out against the NCBI non-redundant protein (nr) database. Transdecoder (v.5.5.0) was used to identify candidate open reading frames and translate genes to proteins for BLASTp analysis against Uniprot reviewed sequence database. Top hit annotations were collated from both BLAST databases with a p-value ≤0.05. Domain analysis was carried out on predicted proteins using Interproscan (v.5.36-75.0) and PFAM domain analysis using hmmscan functions of HMMER (v.3.1) (p-value ≤0.05). Differentially expressed genes also underwent KEGG pathway analysis to identify important biological pathway differences within datasets. BlastKOALA was used to assign KEGG gene K numbers based on matches with known human pathway genes. R (v.3.6.2) packages DESeq2 (v.1.26.0), gage (v.2.36.0) and pathview (v.1.26.0) were used to analyse results using custom R scripts and determine significant differences (p-value ≤0.05). GO term analysis was also carried out on genes annotated in the *F. hepatica* genome using data retrieved from WormBase ParaSite (WBPS14; https://parasite.wormbase.org/index.html) [89]. Data were visualised using GraphPad Prism (v.8) and R (v.3.6.2) packages DESeq2 (v.1.26.0), ggplot2 (v.3.3.1) and upsetR. (v.1.4.0) with custom R scripts.

### Identification and differential expression analysis of miRNAs in in vivo and in vitro maintained F. hepatica

Small RNA fastq files were aligned to a Bowtie index of *F. hepatica* genome PRJEB25283 using miRDeep2 (v 2.0.1.2) [97]. miRNAs were defined using the following criteria; present in at least 2 out of 3 replicates for either *in vitro* or *in vivo* transcriptomes, a minimum of 10 reads mapped to the mature sequence, at least 1 read mapped to the star sequence, a significant randfold value and a minimum miRDeep2 score of 5. miRNA naming was consistent with that presented in Herron et al. [49]. Differential expression of identified miRNAs was carried out using DESeq2 (version 1.28.1), with an adjusted *p*-value of 0.001 for significance. miRNA target prediction was then carried out using miRanda (version 3.3) [98], with thresholds of minimum pairing score of 150 and maximum free energy score of −20. Predicted targets were refined to those differentially expressed in *in vitro* and *in vivo* mRNA transcriptomes and correlating with miRNA expression, i.e. upregulated in *in vivo* miRNA datasets and downregulated in *in vivo* mRNA datasets, for further interpretation. GO terms associated with predicted miRNA targets were retrieved from WormBase ParaSite (WBPS15) and manually interpreted for frequency (number of times GO term was present in datasets) plotted against Log2 fold change from previous transcriptome analysis.

## Acknowledgements

The authors acknowledge the National Centre for the Replacement, Refinement and Reduction of Animals in Research (NC3Rs) grant NC/N001486/1, the Biotechnology and Biological Sciences Research Council (BBSRC) grant BB/T002727/1 and The Department for Education (DfE) and the Department for Agriculture, Environment and Rural Affairs (DAERA) for postgraduate studentship support for DW, EG, NC and RA.

## Supporting information

**S1. File. Transcripts for 21 day old juvenile *Fasciola hepatica* (*in vivo* & *in vitro*)**

**S2. Table. Transcriptome statistics**

**S3. Table. Annotations for (A) all genes, (B) differentially expressed genes and (C) on/off genes**

**S4. Table. (A) Neoblast, (B) cathepsin, (C) metabolomic and (D) neuropeptide markers**

**S5. Fig. (A) Survival of juvenile liver fluke in aerobic and anaerobic conditions.** Percentage survival of juvenile liver fluke across 21 day period post excystment incubated under standard 5% CO2 conditions (red line) and in an anaerobic chamber (purple line). Juveniles showed significant and continuing higher rates of death when maintained in anaerobic chamber after 14 days (2-way ANOVA with Šídák’s multiple comparisons test; **** P<0.001). **(B) Components of N-glycan biosynthesis and processing pathways downregulated in *in vivo* maintained *F. hepatica* juveniles.** Differential expression (log_2_FC) of protein glycosylating genes associated with N-glycan biosynthesis and processing. Genes identified in *F. hepatica* by McVeigh et al. [32] KEGG pathway analysis using R (v.3.6.2), gage (v.2.36.0) and pathview (v.1.26.0) packages identified a significant downregulation of N glycan biosynthesis in *in vivo* maintained juveniles (P≤0.05). Abbreviations; ALG N-glycan precursor synthesis-*Fh-*ALG(−3, −9)=dolichyl-P-Man:Man(5)GlcNAc(2)-PP-dolichol alpha-1,3-mannosyltransferase, *Fh-*ALG5=dolichyl-phosphate beta-glucosyltransferase, *Fh-*ALG7= UDP-N-acetylglucosamine--dolichyl-phosphate N-acetylglucosaminephosphotransferase; Oligosaccharyltransferase complex components-OST48= dolichyl-diphosphooligosaccharide--protein glycosyltransferase subunit, *Fh-*RPN(−1, −2)&Fh-STT3(-A, -B)= dolichyl-diphosphooligosaccharide--protein glycosyltransferase subunit; N-glycan processing-*Fh*-GCNT2=N-acetyllactosaminide beta-1,6-N-acetylglucosaminyl-transferase, *Fh-*FUT8=alpha-(1,6)-fucosyltransferase, B4GALT=beta-1,4-galactosyltransferase, B4GALTNT=Beta--n-acetylgalactosaminyltransferase, EDEM1=ER degradation-enhancing alpha-mannosidase-like protein, UGGT= UDP-glucose:glycoprotein glucosyltransferase, MAN2B1=alpha-mannosidase.

**S6. Table. (A) identified miRNAs (B) novel miRNAs (C) differentially expressed miRNAs (D) miRNA regulated Wnt genes**

## References

1. World Health Organisation. The “Neglected” Neglected Worms. Action Against Worms. 2007. Available from: https://www.who.int/publications/i/item/WHO-HTM-NTD-NZD-2008.2.

2. Spithill T, Smooker P, Coperman D. Fasciola gigantica: epidemiology, control, immunology and molecular biology. In: Dalton J, ed. by. Fasciolosis. 1st ed. Oxworth: Commonwealth Agricultural Bureau International; 1999. p. 465–525.

3. Webb C, Cabada M. Recent developments in the epidemiology, diagnosis, and treatment of Fasciola infection. Current Opinion in Infectious Diseases. 2018;31(5):409–414. PMID: 30113327.

4. Fairweather I, Brennan G, Hanna R, Robinson M, Skuce P. Drug resistance in liver flukes. International Journal for Parasitology: Drugs and Drug Resistance. 2020;12:39–59. PMID: 32179499.

5. Tolan Jr RW. Fascioliasis due to Fasciola hepatica and Fasciola gigantica infection: an update on this ‘neglected’ neglected tropical disease. Laboratory Medicine. 2011 Feb 1;42(2):107–16.

6. Kelley J, Elliott T, Beddoe T, Anderson G, Skuce P, Spithill T. Current Threat of Triclabendazole Resistance in Fasciola hepatica. Trends in Parasitology. 2016;32(6):458–469. PMID: 27049013.

7. Kaplan R. Fasciola hepatica: a review of the economic impact in cattle and considerations for control. Veterinary Therapeutics. 2001;2(1):40–50.

8. Cwiklinski K, Jewhurst H, McVeigh P, Barbour T, Maule A, Tort J et al. Infection by the Helminth Parasite Fasciola hepatica Requires Rapid Regulation of Metabolic, Virulence, and Invasive Factors to Adjust to Its Mammalian Host. Molecular & Cellular Proteomics. 2018;17(4):792–809. PMID: 29321187.

9. John B, Davies D, Williams D, Hodgkinson J. A review of our current understanding of parasite survival in silage and stored forages, with a focus onFasciola hepaticametacercariae. Grass and Forage Science. 2019;74(2):211–217. PMID: 31244499.

10. Gajewska A, Smaga-Kozłowska K, Wiśniewski M. Pathological changes of liver in infection of Fasciola hepatica. Wiad Parazytol. 2005;51(2):115–123. PMID: 16838620.

11. Di Maggio L, Tirloni L, Pinto A, Diedrich J, Yates III J, Benavides U et al. Across intra-mammalian stages of the liver f luke Fasciola hepatica: a proteomic study. Scientific Reports. 2016;6(1). PMID: 27600774.

12. Gil L, Díaz A, Rueda C, Martínez C, Castillo D, Apt W. Fascioliasis hepática humana: resistencia al tratamiento con triclabendazol. Revista médica de Chile. 2014;142(10):1330–1333. PMID: 25601119.

13. Cabada M, Lopez M, Cruz M, Delgado J, Hill V, White A. Treatment Failure after Multiple Courses of Triclabendazole among Patients with Fascioliasis in Cusco, Peru: A Case Series. PLOS Neglected Tropical Diseases. 2016;10(1):e0004361. PMID: 26808543.

14. Branco E, Ruas R, Nuak J, Sarmento A. Treatment failure after multiple courses of triclabendazole in a Portuguese patient with fascioliasis. BMJ Case Reports. 2020;13(3):e232299. PMID: 32193176.

15. McCusker P, McVeigh P, Rathinasamy V, Toet H, McCammick E, O’Connor A et al. Stimulating Neoblast-Like Cell Proliferation in Juvenile Fasciola hepatica Supports Growth and Progression towards the Adult Phenotype In Vitro. PLOS Neglected Tropical Diseases. 2016;10(9):e0004994. PMID: 27622752.

16. Duque-Correa M, Maizels R, Grencis R, Berriman M. Organoids – New Models for Host–Helminth Interactions. Trends in Parasitology. 2020;36(2):170–181. PMID: 31791691.

17. Wang J, Chen R, Collins J. Systematically improved in vitro culture conditions reveal new insights into the reproductive biology of the human parasite Schistosoma mansoni. PLOS Biology. 2019;17(5):e3000254. PMID: 31067225.

18. McVeigh P, McCusker P, Robb E, Wells D, Gardiner E, Mousley A et al. Reasons to Be Nervous about Flukicide Discovery. Trends in Parasitology. 2018;34(3):184–196.PMID: 29269027

19. Young ND, Hall RS, Jex AR, Cantecessi C, Gasser RB. Elucidating the transcriptome of Fasciola hepatica - a key to fundamental and biotechnological discoveries for a neglected parasite. 2010;28(2):222–231. PMID: 20006979

20. Cwiklinski K, Dalton J, Dufresne P, La Course J, Williams D, Hodgkinson J et al. The Fasciola hepatica genome: gene duplication and polymorphism reveals adaptation to the host environment and the capacity for rapid evolution. Genome Biology. 2015;16(1). PMID: 25887684.

21. Radio S, Fontenla S, Solana V, Matos Salim A, Araújo F, Ortiz P et al. Pleiotropic alterations in gene expression in Latin American Fasciola hepatica isolates with different susceptibility to drugs. Parasites & Vectors. 2018;11(1). PMID: 29368659.

22. Cwiklinski K, Robinson MW, Donnelly S, Dalton JP. Complementary transcriptomic and proteomic analyses reveal the cellular and molecular processes that drive growth and development of Fasciola hepatica in the host liver. BMC Genomics. 2021;22(1). PMID: 33430759.

23. Miranda-Miranda E, Cossio-Bayugar R, Aguilar-Díaz H, Narváez-Padilla V, Sachman-Ruíz B, Reynaud E. Transcriptome assembly dataset of anthelmintic response in Fasciola hepatica. Data in Brief. 2021;35:106808. PMID: 33659584.

24. Collins III J, Wang B, Lambrus B, Tharp M, Iyer H, Newmark P. Adult somatic stem cells in the human parasite Schistosoma mansoni. Nature. 2013;494(7438):476-479. PMID: 23426263.

25. Giri B, Li H, Chen Y, Cheng G. Preliminary evaluation of neoblast-like stem cell factor and transcript expression profiles in Schistosoma japonicum. Acta Tropica. 2018;187:57–64. PMID: 30055172.

26. Cwiklinski K, Donnelly S, Drysdale O, Jewhurst H, Smith D, De Marco Verissimo C et al. The cathepsin-like cysteine peptidases of trematodes of the genus Fasciola. Advances in Parasitology. 2019;:113–164. PMID: 31030768.

27. Dalton J, Neill S, Stack C, Collins P, Walshe A, Sekiya M et al. Fasciola hepatica cathepsin L-like proteases: biology, function, and potential in the development of first generation liver fluke vaccines. International Journal for Parasitology. 2003;33(11):1173–1181. PMID: 13678633.

28. Robinson M, Menon R, Donnelly S, Dalton J, Ranganathan S. An Integrated Transcriptomics and Proteomics Analysis of the Secretome of the Helminth Pathogen Fasciola hepatica. Molecular & Cellular Proteomics. 2009;8(8):1891–1907. PMID: 19443417.

29. Grote A, Caffrey C, Rebello K, Smith D, Dalton J, Lustigman S. Cysteine proteases during larval migration and development of helminths in their final host. PLOS Neglected Tropical Diseases. 2018;12(8):e0005919. PMID: 30138448.

30. Wang Q, Da’dara A, Skelly P. The human blood parasite Schistosoma mansoni expresses extracellular tegumental calpains that cleave the blood clotting protein fibronectin. Scientific Reports. 2017;7(1). PMID: 29018227.

31. Tielens A, van Hellemond J. Unusual aspects of metabolism. In: Maule A, Marks N, ed. by. Parasitic flatworms: molecular biology, biochemistry, immunology and physiology. CAB International; 2006. p. 387–408.

32. McVeigh P, Cwiklinski K, Garcia-Campos A, Mulcahy G, O’Neill S, Maule A et al. In silico analyses of protein glycosylating genes in the helminth Fasciola hepatica (liver fluke) predict protein-linked glycan simplicity and reveal temporally-dynamic expression profiles. Scientific Reports. 2018;8(1).

33. Harada Y, Ohkawa Y, Kizuka Y, Taniguchi N. Oligosaccharyltransferase: A Gatekeeper of Health and Tumor Progression. International Journal of Molecular Sciences. 2019;20(23):6074. PMID: 31810196.

34. Hokke C, van Diepen A. Helminth glycomics – glycan repertoires and host-parasite interactions. Molecular and Biochemical Parasitology. 2017;215:47–57. PMID: 27939587.

35. Lin B, Qing X, Liao J, Zhuo K. Role of Protein Glycosylation in Host-Pathogen Interaction. Cells. 2020;9(4):1022. PMID: 32326128.

36. Davis R, Stretton A. Neurotransmitters of Helminths. In: Marr J, Müller M, ed. by. Biochemistry and Molecular Biology of Parasites. Elsevier; 1995.

37. McVeigh P, Kimber M, Novozhilova E, Day T. Neuropeptide signalling systems in flatworms. Parasitology. 2006;131(S1):S41. PMID: 16569292.

38. McVeigh P, Maule A. Flatworm Neurobiology in the Postgenomic Era. In: Byrne JH, ed. by. The Oxford Handbook of Invertebrate Neurobiology. Oxford University Press; 2017.

39. Südhof TC. The synaptic vesicle cycle: a cascade of protein-protein interactions. Nature. 1995;375(6533):645-53. PMID: 7791897.

40. Ribeiro P, El-Shehabi F, Patocka N. Classical transmitters and their receptors in flatworms. Parasitology. 2005;S131:S19–40. PMID: 16569290.

41. Fairweather I, Maule A, Mitchell S, Johnston C, Halton D. *I*mmunocytochemical demonstration of 5-hydroxytryptamine (serotonin) in the nervous system of the liver fluke, Fasciola hepatica (Trematoda, Digenea). Parasitology Research. 1987;73(3): 255–8. PMID: 3295862.

42. Tembe E, Holden-Dye L, Smith S, Jacques P, Walker R. Pharmacological profile of the 5-hydroxytryptamine receptor of Fasciola hepatica body wall muscle. Parasitology. 1993;106(Pt1):67–73. PMID: 8479803.

43. McVeigh P, Mair G, Atkinson L, Ladurner P, Zamanian M, Novozhilova E, Marks N, Day T, Maule A. Discovery of multiple neuropeptide families in the phylum Platyhelminthes. International Journal for Parasitology. 2009;39(11):1243–52. PMID: 19361512.

44. McVeigh P, McCammick E, McCusker P, Wells D, Hodgkinson J, Paterson S, Mousley A, Marks N, Maule A. Profiling G protein-coupled receptors of Fasciola hepatica identifies orphan rhodopsins unique to phylum Platyhelminthes. International Journal for Parasitology: Drugs and Drug Resistance. 2018;8(1):87–103. PMID: 29474932

45. Xu M, Ai L, Fu J, Nisbet A, Liu Q, Chen M et al. Comparative Characterization of MicroRNAs from the Liver Flukes Fasciola gigantica and F. hepatica. PLoS ONE. 2012;7(12):e53387. PMID: 23300925.

46. Fromm B, Trelis M, Hackenberg M, Cantalapiedra F, Bernal D, Marcilla A. The revised microRNA complement of Fasciola hepatica reveals a plethora of overlooked microRNAs and evidence for enrichment of immuno-regulatory microRNAs in extracellular vesicles. International Journal for Parasitology. 2015;45(11):697–702. PMID: 26183562

47. Fontenla S, Dell’Oca N, Smircich P, Tort J, Siles-Lucas M. The miRnome of Fasciola hepatica juveniles endorses the existence of a reduced set of highly divergent micro RNAs in parasitic flatworms. International Journal for Parasitology. 2015;45(14):901–913. PMID: 26432296.

48. Ovchinnikov V, Kashina E, Mordvinov V, Fromm B. EV-transported microRNAs of Schistosoma mansoni and Fasciola hepatica: Potential targets in definitive hosts. Infection, Genetics and Evolution. 2020;85:104528.

49. Herron C, O’Connor A, Robb E, McCammick E, Hill C, Marks N et al. Developmental regulation and functional prediction of microRNAs in an expanded Fasciola hepatica miRNome. 2021;.

50. Kaltz O, Shykoff JA. Local adaptation in host-parasite systems. Heredity. 1998;81(4):361–370.

51. Read AF, Skorping A. The evolution of tissue migration by parasitic nematode larvae. Parasitology. 1995;111(Pt 3):359–71. PMID: 7567104.

52. Orrego-Solano MA, Cangalaya C, Nash T, Guerra-Giraldez C, de Trabajo en Cisticercosis en Peru. Identification of proliferating cells in Taenia solium cyst. Rev Peru Med Exp Salud Publica. 2014;31(4):702–6. PMID: 25597721

53. Baguñà J. The planarian neoblast: the rambling history of its origin and some current black boxes. The International Journal for Developmental Biology. 2012;56(1-3):19–37. PMID: 22252540.

54. Rink JC. Stem cell systems and regeneration in planaria. Developmental Genes and Evolution. 2013;223(1-2):67–84. PMID: 23138344.

55. Koziol U, Rauschendorfer T, Rodríguez LZ, Krohne G, Brehm K. The unique stem cell system of the immortal larva of the human parasite Echinococcus multilocularis. Evodevo. 2014;5(1):10. PMID: 24602211.

56. Collins JJ, Wendt GR, Lyer H, Newmark PA. Stem cell progeny contribute to the schistosome host-parasite interface. Elife. 2016;5:e12473. PMID: 27003592.

57. Collins JN, Collins JJ. Tissue Degeneration following Loss of Schistosoma mansoni cbp1 Is Associated with Increased Stem Cell Proliferation and Parasite Death In Vivo. PLoS Pathogen. 2016;12(11):e1005963. PMID: 27812220

58. Akiba Y, Chida Y, Takahashi T, Ohtomo Y, Sato K, Takahashi K. Persistent hypoglycemia induced by continuous insulin infusion in broiler chickens. British Poultry Science. 1999;40(5):701–5. PMID: 10670686

59. Tielens AG, van den Heuvel JM, van den Bergh. The energy metabolism of Fasciola hepatica during its development in the final host. Molecular and Biochemical Parasitology. 1984;13(3):301–307. PMID: 6527693

60. Alvarado R, To J, Lund ME, Pinar A, Mansell A, Robinson MW, O’Brien BA, Dalton JP, Donnelly S. The immune modulatory peptide FhHDM-1 secreted by the helminth Fasciola hepatica prevents NLRP3 inflammasome activation by inhibiting endolysosomal acidification in macrophages. The FASEB Journal. 2017;31(1):85–95. PMID: 27682204.

61. Martínez-Sernández V, Mezo M, González-Warleta M, Perteguer MJ, Muiño L, Guitián E, Gárate T, Ubeira FM. The MF6p/FhHDM-1 major antigen secreted by the trematode parasite Fasciola hepatica is a heme-binding protein. Journal of Biological Chemistry. 2014;289(3):1441–56. PMID: 24280214.

62. Robinson M, Donnelly S, Dalton JP. Helminth defence molecules-immunomodulators designed by parasites! Frontiers in Microbiology. 2013;4:296. PMID: 24101918

63. Freeze HH, Elbein AD. Glycosylation precursors. In: Varki A, Cummings RD, Esko JD, Stanley P, Hart GW, Aebi M, Darvill AG, Kinoshita T, Packer NH, Prestegard JH, Schnaar RL, Seeberger PH. ed. by. Essentials of Glycobiology. 2009. Cold Spring Harbor Laboratory Press.

64. Fadda M, Hasakiogullari I, Temmerman L, Beets I, Zels S, Schoofs L. Regulation of Feeding and Metabolism by Neuropeptide F and Short Neuropeptide F in Invertebrates. Frontiers in Endocrinology. 2019;10(64). PMID: 30837946.

65. Camacho M, Agnew A. Schistosoma: rate of glucose import is altered by acetylcholine interaction with tegumental acetylcholine receptors and acetylcholinesterase. Experimental Parasitology. 1995;81(4):584–91. PMID: 8543000.

66. Nässel DR, Wegener C. A comparative review of short and long neuropeptide F signaling in invertebrates: Any similarities to vertebrate neuropeptide Y signaling? Peptides. 2011;32(6):1335–55. PMID: 21440021.

67. Wu Q, Zhao Z, Shen P. Regulation of aversion to noxious food by Drosophila neuropeptide Y-and insulin-like systems. Nature Neuroscience. 2005;8(10):1350–5. PMID: 16172603.

68. Chen J, Cheuk I, Shin V, Kwong A. Acetylcholine receptors: Key players in cancer development. Surgical Oncology. 2019;31:46–53. PMID: 31536927

69. Weist R, Flörkemeier T, Roger Y, Franke A, Schwanke K, Zweigerdt R, Martin U, Willbold E, Hoffman A. Differential Expression of Cholinergic System Components in Human Induced Pluripotent Stem Cells, Bone Marrow-Derived Multipotent Stromal Cell, *a*nd Induced Pluripotent Stem Cell-Derived Multipotent Stromal Cells. Stem Cells and Development. 2018;27(3):166–183. PMID: 29205106.

70. Landgraf D, Barth M, Layer PG, Sperling LE. Acetylcholine as a possible signaling molecule in embryonic stem cells: studies on survival, proliferation and death. Chemico-Biological Interactions. 2010. 187(1-3): p. 115–119. PMID: 20223227.

71. Gingle AR. Acetylcholine and carnitine sensitive growth in a Drosophila cell line. Comparative Biochemistry Physiology Part C: Pharmacology and Toxicology. 1985;82(1): p. 235–41. PMID: 2865070.

72. Ruaud AF, Bessereau JL. The P-type ATPase CATP-1 is a novel regulator of C. elegans developmental timing that acts independently of its predicted pump function. Development. 2007;134(5):867–79. PMID: 17251264.

73. Li BW, Rush AC, Weil GJ. Expression of five acetylcholine receptor subunit genes in Brugia malayi adult worms. International Journal for Parasitology: Drugs Drug Resistance. 2015;5(3):100–9. PMID: 26199859.

74. Kreshchenko ND, Functions of flatworm neuropeptides NPF, GYIRF and FMRF in course of pharyngeal regeneration of anterior body fragments of planarian, Girardia tigrina. Acta Biologica Hungarica. 2008; 59:199–207. PMID: 18652393.

75. Collins JJ, Hou X, Romanova EV, Lambrus BG, Miller CM, Saberi A, Sweedler JV, Newmark PA. Genome-wide analyses reveal a role for peptide hormones in planarian germline development. PLoS Biology. 2010;8(10):1000509. PMID: 20967238.

76. Ameku T, Yoshino Y, Texada MJ, Kondo S, Amezawa, Yoshizaki G, Shimida-Niwa Y, Niwa R. Midgut-derived neuropeptide F controls germline stem cell proliferation in a mating-dependent manner. PLoS Biology. 2018;16(9):e2005004. PMID: 30248087.

77. Wang B, Collins JJ, Newmark PA. Functional genomic characterization of neoblast-like stem cells in larval Schistosoma mansoni. Elife. 2013;2:e00768. PMID: 23908765.

78. Saberi A, Jamal A, Beets I, Schoofs L, Newmark P. GPCRs Direct Germline Development and Somatic Gonad Function in Planarians. PLOS Biology. 2016;14(5):e1002457.PMID: 27163480.

79. Shivdasani R. MicroRNAs: regulators of gene expression and cell differentiation. Blood. 2006;108(12):3646–3653.PMID: 16882713.

80. Worringer K, Rand T, Hayashi Y, Sami S, Takahashi K, Tanabe K et al. The let-7/LIN-41 Pathway Regulates Reprogramming to Human Induced Pluripotent Stem Cells by Controlling Expression of Prodifferentiation Genes. Cell Stem Cell. 2014;14(1):40–52.PMID: 24239284.

81. Hunter S, Finnegan E, Zisoulis D, Lovci M, Melnik-Martinez K, Yeo G et al. Functional Genomic Analysis of the let-7 Regulatory Network in Caenorhabditis elegans. PLoS Genetics. 2013;9(3):e1003353.PMID: 23516374.

82. Makeyev E, Zhang J, Carrasco M, Maniatis T. The MicroRNA miR-124 Promotes Neuronal Differentiation by Triggering Brain-Specific Alternative Pre-mRNA Splicing. Molecular Cell. 2007;27(3):435–448.PMID: 17679093.

83. Li K, Pang J, Ching A, Wong C, Kong X, Wang Y et al. miR-124 is frequently down-regulated in medulloblastoma and is a negative regulator of SLC16A1. Human Pathology. 2009;40(9):1234–1243.PMID: 19427019.

84. Song J, Nigam P, Tektas S, Selva E. microRNA regulation of Wnt signaling pathways in development and disease. Cellular Signalling. 2015;27(7):1380–1391.

85. McVeigh P, McCammick E, McCusker P, Morphew R, Mousley A, Abidi A et al. RNAi Dynamics in Juvenile Fasciola spp. Liver Flukes Reveals the Persistence of Gene Silencing In Vitro. PLoS Neglected Tropical Diseases. 2014;8(9):e3185. PMID: 25254508.

86. Martin M. Cutadapt removes adapter sequences from high-throughput sequencing reads. EMBnetjournal. 2011;17(1):10.

87. Joshi N. Sickle: A sliding-window, adaptive, quality-based trimming tool for FastQ files (version 1.33). 2011. Available from: https://github.com/najoshi/sickle.

88. Andrews S. FastQC: A quality control tool for high throughput sequence data. 2010. Available from: http://www.bioinformatics.babraham.ac.uk/projects/fastqc/.

89. Howe K, Bolt B, Shafie M, Kersey P, Berriman M. WormBase ParaSite − a comprehensive resource for helminth genomics. Molecular and Biochemical Parasitology. 2017;215:2–10. PMCID: PMC5486357.

90. Trapnell C, Williams B, Pertea G, Mortazavi A, Kwan G, van Baren M et al. Transcript assembly and quantification by RNA-Seq reveals unannotated transcripts and isoform switching during cell differentiation. Nature Biotechnology. 2010;28(5):511–515. PMID: 20436464.

91. Kim D, Langmead B, Salzberg S. HISAT: a fast spliced aligner with low memory requirements. Nature Methods. 2015;12(4):357–360. PMID: 25751142.

92. Pertea M, Pertea G, Antonescu C, Chang T, Mendell J, Salzberg S. StringTie enables improved reconstruction of a transcriptome from RNA-seq reads. Nature Biotechnology. 2015;33(3):290–295. PMID: 25690850.

93. Pertea M, Kim D, Pertea G, Leek J, Salzberg S. Transcript-level expression analysis of RNA-seq experiments with HISAT, StringTie and Ballgown. Nature Protocols. 2016;11(9):1650–1667. PMID: 27560171.

94. Love M, Huber W, Anders S. Moderated estimation of fold change and dispersion for RNA-seq data with DESeq2. Genome Biology. 2014;15(12). PMID: 25516281.

95. R Core Team. R: A language and environment for statistical computing. R Foundation for Statistical Computing, Vienna, Austria. 2018. Available from: http://www.R-project.org.

96. Love MI, Anders S, Huber W. Analyzing RNA-seq data with DESeq2. 2020. Available from: http://bioconductor.org/packages/devel/bioc/vignettes/DESeq2/inst/doc/DESeq2.html

97. Friedländer M, Mackowiak S, Li N, Chen W, Rajewsky N. miRDeep2 accurately identifies known and hundreds of novel microRNA genes in seven animal clades. Nucleic Acids Research. 2011;40(1):37–52.PMID: 21911355.

98. Betel D, Koppal A, Agius P, Sander C, Leslie C. Comprehensive modeling of microRNA targets predicts functional non-conserved and non-canonical sites. Genome Biology. 2010;11(8):R90.PMID: 20799968.

